# Selection scans and downstream analysis with selscan

**DOI:** 10.1101/2025.10.10.681670

**Authors:** Amatur Rahman, T. Quinn Smith, Zachary A. Szpiech

## Abstract

Summary statistics based on Extended Haplotype Homozygosity (EHH) are widely used for inferring positive selection in genomes as a result of their ease of use, computational efficiency, and interpretability. These various summary statistics can be applied to single populations or to pairs of populations, can be used with a genetic recombination map or without, and can be applied to phased or unphased data. Although these statistics are straightforward to compute, there lacks clear descriptions on how they relate to one another, how they should be used, and how their resulting outputs should be interpreted. Here, we provide a comprehensive introduction to selection statistics as they are implemented in the widely used software, `selscan`. In addition to this detailed guide, we implement enhanced normalization procedures and support for gene-based analyses, enabling users to translate selection signals captured by these statistics into gene-level interpretations using BED annotation files, facilitating biologically meaningful insights. We demonstrate the behavior of such statistics on simulated data and highlight best practices by performing an example downstream analysis on data from the 1000 Genomes Project using new features in `selscan v3.0`. We hope these guidelines will foster reproducibility in the evolutionary genomics community. Precompiled executables and source code for `selscan v3.0` can be found at https://github.com/szpiech/selscan.

## 1 Introduction

Among evolutionary biologists, there is great interest in identifying the genomic basis of adaptations. Pinpointing putatively adaptive mutations, especially in relation to known selective pressures and with functional validation, can provide insights into the biological mechanisms underlying phenotypic innovations. In humans, understanding the biological basis of adaptation can give a deeper understanding of our evolutionary history and can reveal important information about the relationship between genes, adaptation, and disease (Scheinfeldt and Tishkoff, 2013; Fu and Akey, 2013; Benton et al., 2021).

Given this importance, many statistics have been developed to make inferences about positive selection in the genome across different time scales (Booker et al., 2017; Pavlidis and Alachiotis, 2017). For detecting recent and strong positive selection, common methodologies typically make use of expected distortions in either the site frequency spectrum or haplotype patterns (Szpiech and Hernandez, 2016). When a strongly adaptive allele arises, it will sweep to high frequency on a timescale faster than recombination or mutation occur, bringing linked variation to high frequency as well. This results in a region of low genetic diversity, long haplotypes, and an excess of high and low frequency alleles. Although more complicated model-based (Nielsen et al., 2005; DeGiorgio et al., 2016; Harris and DeGiorgio, 2020; DeGiorgio and Szpiech, 2022; Stern et al., 2019; Vaughn and Nielsen, 2024; Hejase et al., 2022) and machine learning (Kern and Schrider, 2018; Sugden et al., 2018; Amin et al., 2024; Arnab et al., 2025) methods have proven useful, especially for inferring specific parameters of a sweep, summary-statistic-based methods remain widely used as a result of their ease of use, computational efficiency, and interpretability.

One particularly important class of summary statistics for inferring positive selection are those based on extended haplotype homozygosity (Sabeti et al., 2002), which are designed to capture the signal of long high frequency haplotypes in the vicinity of a sweep that is either ongoing or recently completed. EHH-based summary statistics include iHS (Voight et al., 2006), nSL (Ferrer-Admetlla et al., 2014), XP-EHH (Sabeti et al., 2007), and XP-nSL (Szpiech et al., 2021), and although originally formulated for use on phased haplotype data, they have also been extended to work on unphased data as well (Klassmann and Gautier, 2022; Szpiech, 2024). These statistics are also frequently included as important components in more advanced machine learning approaches (Amin et al., 2024; Arnab et al., 2025). Given their broad adoption for use in selection inference analyses, they have been implemented in several widely used software programs including `selscan` (Szpiech and Hernandez, 2014; Szpiech, 2024; Rahman et al., 2025), rehh (Gautier and Vitalis, 2012; Gautier et al., 2017), and hapbin (Maclean et al., 2015). Currently, selscan is the leading implementation in terms of computational efficiency and memory usage (Rahman et al., 2025).

In this work, we introduce the definitions of these common EHH-based statistics and their interpretations. We also provide an overview of how to use `selscan v3.0` for genome-wide selection inference and downstream analysis, while highlighting new features such as improved normalization and gene-based analysis support that allow users to map selection signals to genes using BED annotation files for interpretable functional insights.

## 2 Materials and Methods

First, we introduce the main statistics (Sections 2.1, 2.2, 2.3, and 2.4) and provide the exact commands used on simulated data. These commands match those used to produce the figures and results in the Results section, aiming to provide a quick start for users interested in performing selection scan analyses. We also highlight the importance of downstream analysis and present both its application and the relevant commands (Section 2.5). We describe our simulations and dataset preprocessing methods in Section 2.7.

### 2.1 EHH

#### 2.1.1 Definition for phased data

Extended Haplotype Homozygosity (EHH) is the probability that two chromosomes randomly chosen from those carrying a particular “core” allele at a given locus are identical by descent across all markers from that core locus to another position (Sabeti et al., 2002). Consider a sample of *n* chromosomes. Define 𝒞 to be the set of all possible distinct haplotypes at a locus of interest, which is called *x*_0_. In addition, define 𝒞 (*x*_*i*_) to denote the set of all possible distinct haplotypes extending from the locus *x*_0_ to the *i*^*th*^ marker either upstream or downstream from *x*_0_. If *x*_0_ is a biallelic locus, then let 0 represent the ancestral allele and 1 represent the derived allele. Therefore, 𝒞 (*x*_0_) := {0, 1}. If *x*_1_ is an immediately adjacent locus, then the set of all possible haplotypes becomes 𝒞 (*x*_1_) := {11, 10, 00, 01}.

EHH between *x*_0_ and *x*_1_ of the entire sample (Sabeti et al., 2002; Szpiech and Hernandez, 2014) is calculated as

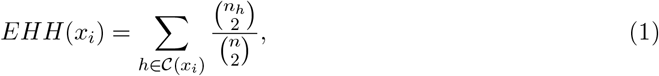

where *n*_*h*_ is the number of observed haplotypes of type *h* ∈ 𝒞 (*x*_*i*_).

It is advantageous to calculate the haplotype homozygosity of a sub-sample of chromosomes all carrying a ‘core’ allele at locus *x*_0_. Define ℌ_*c*_(*x*_*i*_) to be a partition of 𝒞 (*x*_*i*_) containing all unique haplotypes carrying the core allele, *c* ∈ 𝒞, at *x*_0_ and extending to marker *x*_*i*_ as

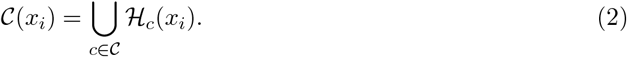

ℌ_1_(*x*_1_) := {11, 10} and ℌ_0_(*x*_1_) := {00, 01} when the core allele is chosen as the derived and ancestral allele at *x*_0_, respectively.

Finally, define EHH of haplotypes containing the core allele, *c*, to a locus *x*_*i*_ as

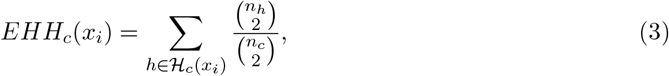

where *n*_*h*_ is the number of chromosomes with haplotype *h* ∈ ℌ_*c*_(*x*_*i*_) and *n*_*c*_ is the number of chromosomes carrying the core allele (*c* ∈ 𝒞).

#### 2.1.2 Definition for unphased data

EHH can be adapted to unphased genotypes of diploid individuals by encoding each locus with the number of observed derived alleles: 0, 1, 2 corresponding to homozygous ancestral, heterozygous, and homozygous derived, respectively. Let 𝒞 := {0, 1, 2} denote the set of all possible genotypes at locus *x*_0_. Define the set of all unique haplotypes extending from site *x*_0_ to site *x*_*i*_ as 𝒞 (*x*_*i*_), where *x*_*i*_ is either upstream or downstream of *x*_0_. For example, if *x*_1_, the site adjacent to *x*_0_, then 𝒞 (*x*_1_) := {00, 01, 02, 10, 11, 12, 20, 21, 22}. With these updated definitions, EHH can be computed for a set of ‘multi-locus genotypes’ using Equation 1.

Similarly, the above reasoning can be applied when defining EHH for a ‘core’ allele. To compute the EHH of a subset of observed haplotypes that all contain the same ‘core’ genotype, let ℌ_*c*_(*x*_*i*_) be the partition of 𝒞 (*x*_*i*_) containing genotype *c* ∈ 𝒞 at *x*_0_. For example, choosing the derived homozygous genotype (*c* = 2) as the core allele, ℌ_2_(*x*_1_) := {20, 21, 22}. Thus, EHH can be computed for all individuals carrying a given genotype at site *x*_0_ extending out to site *x*_*i*_ using Equation 3

Finally, define cEHH as the complement EHH of a sample of haplotypes. cEHH is the EHH of all haplotypes without the ‘core’ genotype at *x*_0_.

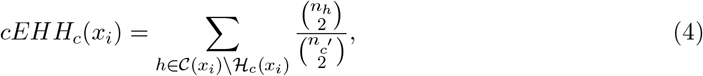

where *n*_*c*_*′* is the number of observed haplotypes with a core genotype of not *c*.

#### 2.1.3 Interpretation of EHH

EHH measures the decay of haplotype homozygosity in iteratively larger windows extending from a core locus. It is computed from among all haplotypes in the sample (Equation 1) or from among only haplotypes containing a particular core allele (Equation 3). Under neutrality, haplotype homozygosity is expected to decay quickly as the window grows larger—reflecting the underlying diversity of the haplotypes in the sample. However, haplotype homozygosity will be maintained at high levels in the vicinity of a recent or ongoing sweep. In this case, EHH of all samples (Equation 1) is expected to decay slowly as window size increases, as the sweeping haplotype will dominate the homozygosity calculation. If computing EHH from among only haplotypes containing a particular core allele (Equation 3), the expected pattern depends on whether the core allele is (or is linked to) the adaptive allele. When the core allele is (or is linked to) the adaptive allele, EHH computed from among only the sweeping haplotypes will have a slow decay as window size increases. On the other hand, the EHH computed from among only non-sweeping haplotypes will have a fast decay as window size increases, similar to the expectation under neutrality. See Section 3.1.2 for an illustration.

#### 2.1.4 Example Commands

To calculate EHH at a marker *rs*1 on phased data:

~~~
selscan --ehh rs1 --vcf input.vcf
~~~

Assuming the input VCF file contains one chromosome, we can calculate EHH at a specific position. For example, at 5MB:

~~~
selscan --ehh 5000000 --vcf input.vcf
~~~

Suppose we have a file with a list of three sites, *genetic*.*map*:

| chr# | ID# | Genetic Position | Physical Position |
| --- | --- | --- | --- |
| 1 | rs1 | 0.09 | 1000010 |
| 1 | rs2 | 0.10 | 1000013 |
| 1 | rs3 | 0.12 | 2000000 |

Then, to run EHH at each of the three loci:

~~~
selscan --ehh 1000010,1000013,2000000 --vcf input.vcf --map genetic.map
~~~

To run on unphased data, we add the `--unphased` flag.

~~~
selscan --ehh rs1 --vcf input.vcf --map genetic.map --unphased
~~~

### 2.2 iHS and nSL

#### 2.2.1 Definition for phased data

The decay of haplotype homozygosity tracked by *EHH* and *EHH*_*c*_ provides useful biological information for identifying sweeps at individual loci of interest, but these statistics are not well-suited for genome-wide scans. This motivated the development of iHS, which uses Equation 3 to summarize the decay of ancestral and derived haplotype homozygosity into a single score. iHS is calculated at a site by first calculating the integrated haplotype homozygosity (iHH) for the ancestral (0) and derived (1) haplotypes (𝒞 := {0, 1}) via trapezoidal quadrature.

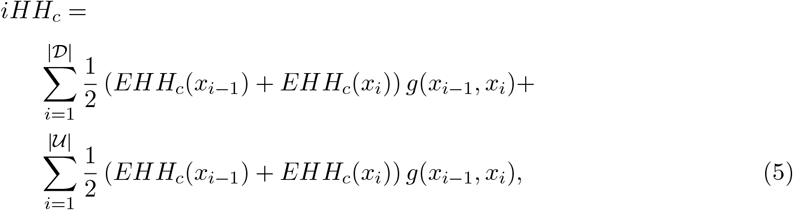

𝒟 is the set of downstream loci, such that, if *x*_*i*_ ∈ 𝒟, then *x*_*i*_ is the *i*^*th*^ closest downstream locus from *x*_0_. Similarly define 𝒰 to be the set of upstream markers. Let *g*(*x*_*i*−1_, *x*_*i*_) be the genetic or physical distance between two loci. The unstandardized iHS is calculated as

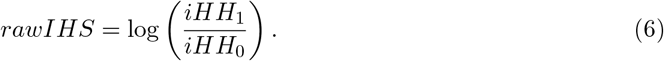

This ensures that when the derived allele is under selection and *iHH*_1_ *> iHH*_0_, the ratio exceeds 1, the log is positive, reflecting the typical case where a positive statistic indicates selection on the derived allele. Note, this definition is different from that in (Voight et al., 2006), where the roles of *iHH*_1_ and *iHH*_0_ are swapped and a natural logarithm is used.

A related statistic, nSL, was introduced by (Ferrer-Admetlla et al., 2014). Although Ferrer- Admetlla et al. define nSL in terms of the mean number of sites shared among all pairwise haplotypes in the vicinity of a query site, they also prove a reformulation in terms of haplotype homozygosity. Using Equation 5 above, the essential difference between nSL and iHS is that the distance function for nSL is given by *g*(*x*_*i*_, *x*_*j*_) = |*j* − *i*|, which simply counts the number of observed segregating sites between *x*_*i*_ and *x*_*j*_.

#### 2.2.2 Definition for unphased data

iHS and nSL can be adapted to unphased genotypes by using the definitions in Section 2.1.2 and defining the homozygous ancestral and the homozygous derived genotypes as *c* = 0 and *c* = 2, respectively. The integrated haplotype homozygosity (iHH) can be calculated for each genotype using Equation 5, and the complement integrated haplotype homozygosity (ciHH) can be calculated for both homozygous core genotypes as

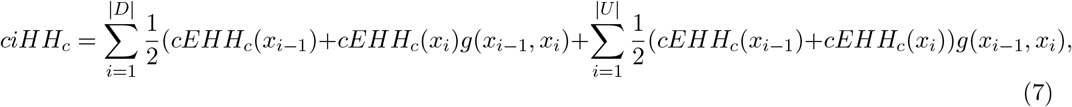

where *g*(*x*_*i*−1_, *x*_*i*_) is the genetic or physical distance between two loci.

The unstandardized unphased iHS is calculated as

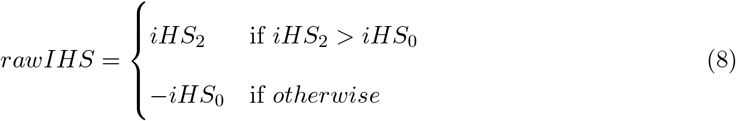

where *iHS*_2_ = *log*((*iHH*_2_)*/*(*ciHH*_2_)) and *iHS*_0_ = *log*((*iHH*_0_)*/*(*ciHH*_0_)). Unstandardized unphased nSL is computed similarly but with *g*(*x*_*i*_, *x*_*j*_) = |*j* − *i*|.

#### 2.2.3 Interpretation of iHS and nSL

iHS/nSL are intended to efficiently contrast the decay of haplotype homozygosity computed from among haplotypes containing the ancestral and derived alleles at a locus of interest. From Equation 6, when the log-ratio takes an extreme positive value, this suggests haplotypes carrying the derived allele are unusually long and low-diversity compared to haplotypes carrying the ancestral allele. When the log-ratio takes an extreme negative value, this suggests haplotypes carrying the ancestral allele are unusually long and low-diversity compared to haplotypes carrying the derived allele. It is tempting, in this case, to think that only extreme positive values are of interest, assuming that an adaptive allele is necessarily derived. However, ancestral alleles in linkage disequilibrium with the adaptive allele also provide information in the vicinity of a sweep, and in some data sets the adaptive allele may not even be observed. Therefore analyses using iHS and nSL are typically concerned with extreme absolute scores. See Section 3.1.2 for an illustration.

#### 2.2.4 Example Commands

To calculate iHS on phased data, `selscan` requires genetic data (e.g., VCF) and a genetic map file containing both physical distances and genetic distance.

~~~
selscan --ihs --vcf input.vcf --map genetic.map
~~~

To calculate iHS on unphased data, add the `--unphased` flag.

~~~
selscan --ihs --vcf input.vcf --map genetic.map --unphased
~~~

When a genetic map is unavailable for iHS, the `--pmap` flag can be used so that selscan approximates genetic distance using physical distance assuming constant recombination. Note that for legacy input formats such as HAP and THAP, a map file is still required even when `--pmap` is specified. In this case, the genetic distance column is ignored, and only the variant ID and physical position columns are used.

~~~
selscan --ihs --vcf input.vcf --pmap
~~~

To calculate nSL on phased data, `selscan` requires only genetic data (e.g., VCF). It uses physical postions included in it. .

~~~
selscan --nsl --vcf input.vcf
~~~

To calculate nSL on unphased data, add the `--unphased` flag.

~~~
selscan --nsl --vcf input.vcf --unphased
~~~

### 2.1 XPEHH and XPnSL

#### 2.3.1 Definition for phased data

XPEHH (Sabeti et al., 2007) and XPnSL (Szpiech et al., 2021) were introduced as two-population extensions to iHS and nSL, respectively, intended to identify local adaptation. For these statistics, instead of comparing the EHH between haplotypes with the ancestral and derived alleles (i.e., by computing *EHH*_*c*_ for each allele type), they compare the EHH of all haplotypes in one population to another population. Consider two populations, *A* and *B*, and a locus *x*_0_. First, iHH is calculated for each population by integrating the EHH of all samples (Equation 1) in each population.

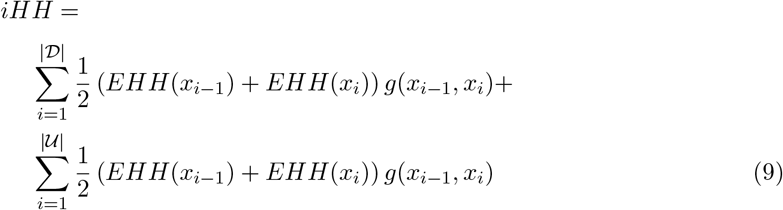

Let *iHH*_*A*_ and *iHH*_*B*_ be the iHH for populations *A* and *B*, respectively. Then the unstandardized XPEHH is

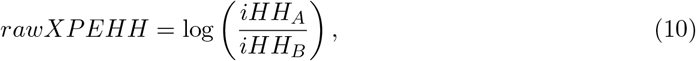

XPnSL is calculated similarly, but with *g*(*x*_*i*−1_, *x*_*i*_) = |*j* − *i*|.

#### 2.3.2 Definition for unphased data

Unphased XPEHH and XPnSL can be calculated using the unphased definitions given in Section 2.1.2 and Equation 9, which defines unphased iHH for the entire population. The distance measure is either centimorgans or base pairs for XPEHH (Sabeti et al., 2007), or the number of observed sites for XPnSL (Szpiech et al., 2021). Both XP statistics between population A and B are computed as

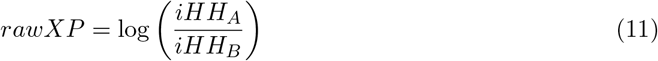

#### 2.3.3 Interpretation of XPEHH and XPnSL

XPEHH/XPnSL are intended to efficiently contrast the decay of haplotype homozygosity of a sample of haplotypes from one population to a sample of haplotypes from a closely related population at a locus of interest. From Equation 10, when the log-ratio takes an extreme positive value, this suggests haplotypes from the “A” population are unusually long and low-diversity compared to haplotypes from the “B” population. When the log-ratio takes an extreme negative value, this suggests haplotypes from the “B” population are unusually long and low-diversity compared to haplotypes from the “A”. In this case, extreme positive values and extreme negative values are of interest, but they should be analyzed separately, as they suggest possible sweeps in one population or the other. See Section 3.1.3 for an illustration. Importantly, in the case where a sweep is occurring in both populations at the same locus, large values of iHH in the log-ratio would cancel out and the result is not distinguishable from neutrality.

#### 2.3.4 Example Commands

To calculate XPEHH on phased data, selscan requires three input files, genetic data (e.g., VCF) for a reference population, genetic data for an alternate population, and a genetic map. We use *alt*.*vcf* to refer to samples from population “A” and *ref*.*vcf* to refer to samples from population “B” in Equations 10 and 11

~~~
selscan --xpehh --vcf-ref ref.vcf --vcf alt.vcf --map genetic.map
~~~

To calculate XPEHH on on unphased data, add the `--unphased` flag.

~~~
selscan --xpehh --vcf-ref ref.vcf --vcf alt.vcf --unphased --map genetic.map
~~~

As with iHS, in the absence of a genetic map, the `--pmap` flag can also be used for XPEHH. To calculate XPnSL on phased data, only two input files are required, genetic data (e.g., VCF) for a reference population, and genetic data for an alternate population.

~~~
selscan --xpnsl --vcf-ref ref.vcf --vcf alt.vcf
~~~

To calculate XPnSL on on unphased data, add the `--unphased` flag.

~~~
selscan --xpnsl --vcf-ref ref.vcf --vcf alt.vcf --unphased
~~~

### 2.4 Integrated Haplotype Homozygosity Pooled (iHH12)

The iHH12 (Torres et al., 2018) statistic is adapted from the H12 statistic (Garud et al., 2015) and is similar to iHH (Equation 9). However iHH12 integrates over EHH12, which is given by

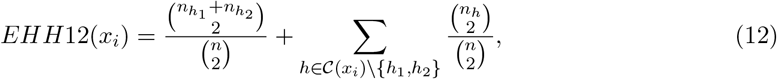

where *h*_*i*_ is the *i*^*th*^ most frequent haplotype in the sample and 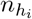 is the number of observed *h*_*i*_ haplotypes. iHH12 is computed with Equation 9 but using *EHH*12 in place of *EHH*.

#### 2.4.1 Interpretation of iHH12

iHH12 is intended to efficiently summarize the decay in haplotype homozygosity among all samples in a single population, with a particular emphasis on detecting soft sweeps. On the assumption that soft sweeps will likely have more than one haplotype sweeping to high frequency, iHH12 combines the counts of the two most frequent haplotypes into a single frequency class, thereby recovering homozygosity signal that would otherwise be lost by partitioning the homozygosity contribution between two classes. When iHH12 values are large, this suggests a collection of long haplotypes of low diversity, and may be indicative of a sweep in the vicinity of that locus. See Section 3.1.2 for an illustration.

#### 2.4.2 Example Commands

To calculate iHH12 on phased data, `selscan` requires genetic data (e.g., VCF) and a genetic map file.

~~~
selscan --ihh12 --vcf input.vcf --map genetic.map
~~~

Currently, there is no unphased version of iHH12.

### 2.5 Downstream analysis

`selscan`‘s main functions compute unstandardized EHH-based statistics. However, normalization is necessary to draw meaningful conclusions, and other analyses, such as outlier detection and gene-based scoring, are commonly performed but require special care in executing. `selscan v3.0` now includes a subcommand, `norm`, to facilitate downstream analysis of selscan results. In the following, we describe how `selscan`‘s `norm` subcommand performs normalization, outlier detection, and gene scoring for potential selection signals.

#### 2.5.1 Normalization

Normalization transforms the raw statistics to an approximate Standard Normal distribution, allowing them to be used for further statistical analysis and to be comparable between studies. For iHS and nSL (phased or unphased) this is a particularly important step, as the raw statistics are correlated with derived allele frequency, and therefore normalization is performed within frequency bins along the genome (see Section 3.1.1 for an illustration). These statistics are therefore normalized by

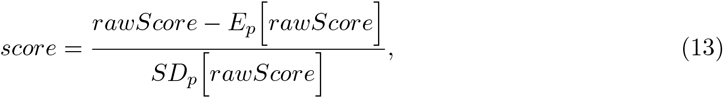

where *E*_*p*_ [*rawScore*] and *SD*_*p*_ [*rawScore*] are the expectation and standard deviation in frequency bin *p*, respectively, of raw scores.

It is recommended to normalize all data together, therefore `selscan norm` accepts multiple `selscan` output files. Normalization of iHS or nSL output using 100 frequency bins can be performed with the following command.

~~~
selscan norm [--ihs|--nsl] --files *.out --bins 100
~~~

XPEHH and XPnSL scores (phased or unphased) and iHH12 scores are not correlated with derived allele frequency (see Section 3.1.1) and are therefore normalized by

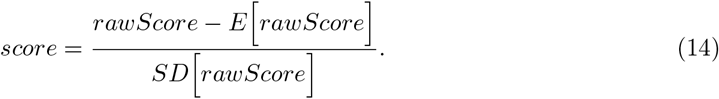

where *E* [*rawScore*] and *SD* [*rawScore*] are the expectation and standard deviation, respectively, of raw scores across the genome.

Normalization of XPEHH, XPnSL, or iHH12 output can be performed with the following command.

~~~
selscan norm [--ihh12|--xpehh|--xpnsl] --files *.out
~~~

##### Speeding Up Normalization

Sometimes it is desirable to normalize a set of data using a set of reference data. This is most commonly encountered when using neutral simulations as a background, either for normalizing empirical data or for normalizing individual simulation replicates. Repeatedly normalizing against the same reference data can be time-consuming. `selscan norm` accommodates this by logging the normalization information, which can then be passed on the command line to be used for normalizing other output.

~~~
selscan norm [--ihs|--nsl|--ihh12|--xpehh|--xpnsl] --log neutral.log \ 
--files neutral_rep*.out
selscan norm [--ihs|--nsl|--ihh12|--xpehh|--xpnsl] --log-input neutral.log \ 
--files sweep_rep*.out
~~~

#### 2.5.2 Window-based outlier detection

While individual extreme scores are suggestive of a sweep, it has been shown that searching for clusters of extreme scores increases power (Voight et al., 2006; Szpiech et al., 2021). To facilitate this type of analysis, `selscan norm` can perform window-based outlier detection, by searching for non-overlapping windows in the genome that contain a high percentage of “extreme” scores. The following command normalizes raw `selscan` output and performs an outlier analysis in nonoverlapping 100kb windows.

~~~
selscan norm [--ihs|--nsl|--ihh12|--xpehh|--xpnsl] --files *.out --bp-win \ 
--winsize 100000
~~~

If we desire to do normalization and windowing separately we replace the command with the following:

~~~
selscan norm [--ihs|--nsl|--ihh12|--xpehh|--xpnsl] --files *.out
selscan norm [--ihs|--nsl|--ihh12|--xpehh|--xpnsl] --norm-files *.out.norm --bp-win \ 
--winsize 100000
~~~

For iHS, nSL, and iHH12, this will compute the percentage of scores within each window for which |*score*| *> C*, for positive *C*. For XPEHH and XPnSL, this will make two calculations: the percentage of scores within each window for which *score > C* and the percentage of scores for which *score <* −*C*. This allows windows to be identified which indicate an enrichment in either population. By default *C* = 2 as suggested in previous studies (Voight et al., 2006; Szpiech et al., 2021), however this value can be changed with --crit-val. Windows are binned into quantiles by number of scores (default 10 bins, use `--qbins` to change) then annotated as being in the top 1% or top 5% within each quantile.

The top 1% windows are typically chosen as outliers when identifying candidate regions under selection. While strong sweep candidates often appear in the top 1%, signals at loci that are weaker may only become visible when using a broader threshold, such as the top 5%. `selscan norm` also provides an option to examine finer-grained percentiles by setting `--fine-percentile`. This allows users to annotate each window with an integer percentile from 1 to 100 (i.e., 1%, 2%, 3%, …, 100%).

#### 2.5.3 Gene-based analyses

It is often desirable to examine which genes (or other genomic features) intersect with the outlier windows identified with`--bp-win` and to compute the maximum observed score per gene. The latter information may be used to rank genes or to perform enrichment tests. `selscan norm` can intersect and annotate windows and compute maximum observed scores per gene when given a BED file.

If `all chr.bed` contains a list of intervals defining gene regions and IDs (could be from multiple chromosomes) and `chr*.out.norm.100kb.windows` are the output files from `selscan norm --bp-win`, the following command can be executed.

~~~
selscan norm [--ihs|--nsl|--ihh12|--xpehh|--xpnsl] \
--win-files chr*.out.norm.100kb.windows --gene-bed all_chr.bed
~~~

This command generates one output file per window file, using gene annotations from the BED file. For example, if `chr1.ihs.out.norm.100kb.windows` is in the list of files, it generates `chr1.ihs.out.norm.100kb.windows.ann`, which contains the windows annotated with overlapping gene names.

A complementary way of viewing gene-based results is to do it via gene table or gene-aggregated SNP scores: The gene table summarizes per-gene scores, allowing ranking or comparison across genes. Although we provide maximum score corrected for length, still caution is advised when interpreting these ranking, as they can be influenced by variability in SNP density, recombination rate, and other factors. These gene-level scores can also be used for enrichment tests. In any case, a single gene table is produced containing all genes and all scores, which can be used for other downstream analyses.

To get SNP level gene table from normalized outputs

~~~
selscan norm [--ihs|--nsl|--ihh12|--xpehh|--xpnsl] \
--files chr*.[ihs|nsl|ihh12|xpehh|xpnsl].out --gene-bed all_chr.bed
~~~

If say, we use iHS scores, this outputs `all chr.ihs.genetable`, which summarizes results at the gene level. For single population statistics, the gene table has nine columns: *chromosome, gene name, start, end, gene length, number of overlapped SNPs, fraction of overlapped SNPs above critical threshold, maximum score*, and *maximum score corrected for gene length*.

The final column is given by the residuals after regressing the score onto gene length. This is an important adjustment as maximum observed score is correlated with gene length (see Section 3.2.1), and gene length is correlated with many biologically interesting features (Lopes et al., 2021). To correct for potential dependence of scores on gene length, we perform a linear regression of the per-gene score on the logarithm of gene length:

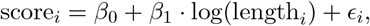

where score_*i*_ is the score assigned to gene *i*, length_*i*_ is the length of the gene, *β*_0_ and *β*_1_ are the regression coefficients, and *ϵ*_*i*_ is the residual. The residuals *ϵ*_*i*_ are returned as length-corrected scores, allowing fair comparison of scores across genes of different lengths.

### 2.6 Other practical considerations

#### 2.6.1 Applying selscan to non-ideal datasets

The performance of EHH-based statistics depends strongly on data quality, yet studies in non-model organisms often rely on imperfect datasets (e.g., sparse SNP density, uncertain genetic maps, missing genotypes, or small sample sizes). These limitations directly affect EHH estimation and can lead to unstable scores or many omitted loci. Simulation studies confirm that SNP density, sample size, and inter-marker distance strongly influence the power of iHS- and XP-EHH-based methods; for example, (Ma et al., 2015) reported that densities of at least ∼1 SNP/kb provide reasonable power in many settings.

In this section, we focus on practical issues that affect EHH estimation—SNP density, map quality, window size, normalization, and missingness—and describe how selscan parameters can be tuned to stabilize scores under imperfect data. Importantly, `selscan` provides window-based summaries (e.g., SNP counts, score distributions, and threshold-based fractions) that help diagnose where data are insufficient and improve interpretability through window aggregation. After preprocessing, selecting statistics that match sweep completeness, reference population availability, and phasing quality—and combining evidence across complementary signals with downstream validation—provides a practical way to make the most of imperfect data.

##### Missing output

When `selscan` cannot compute a statistic at a locus, it typically omits that site from the output (i.e., no score is reported) and may emit a warning. At the SNP level, omitted sites indicate loci where the statistic is undefined or cannot be estimated reliably due to data limitations (e.g., missing genotypes, sparse SNP density, contig/chromosome boundaries, or unreliable genetic maps) or methodological constraints (e.g., rare alleles, rapid EHH decay that fails integration cutoffs, or large inter-marker gaps). Near chromosome boundaries, insufficient flanking markers can also cause omission; the `--trunc-ok` option allows `selscan` to report truncated integration results instead. Genetic map artifacts (e.g., heavy interpolation or collapsed genetic distances resulting in effectively zero integration distance) can further increase omission. Finally, iHS, nSL, and iHH12 are not defined at monomorphic sites, so such loci are skipped. If users observe more omitted sites than expected in a new non-model dataset, it may be helpful to inspect the distribution of missingness and, if appropriate for the data, experiment with tuning `selscan` parameters—for example by adjusting the minor allele frequency cutoff, gap-handling and integration settings such as `--max-gap, --max-distance` (or `--max-distance-nsl`), and the EHH decay cutoff. Masking problematic genomic regions where genetic distances are unreliable can also help prevent systematic missingness. However, missing data is not particularly detrimental to downstream analyses, since we rely on window-based aggregation rather than interpreting individual loci. Within this framework, occasional missing values in a window do not bias the overall inference, as illustrated in the next paragraph.

##### Window-based summaries and tuning for non-ideal datasets

Because locus-level EHH statistics can be noisy and missingness can be substantial in non-model datasets, window-based aggregation provides a practical way to improve interpretability. `selscan` reports window-based summaries that include the number of SNPs contributing to each window, the fraction of SNPs exceeding a critical threshold, and a percentile rank based on thresholds derived from the normalization data. These diagnostics help identify windows with insufficient information (e.g., too few valid SNPs), which are typically flagged as -1 in `selscan`‘s window reports. Window size should be chosen to balance noise reduction with resolution: the `selscan` default is 100 kb, but low-density datasets (e.g., cattle arrays) may require larger windows (e.g., 1 Mb for cattle as in (Randhawa et al., 2014)), whereas high-density datasets (e.g., *Zea mays*) may support smaller windows (e.g., 50 kb as in (Gage et al., 2018)). A practical guideline is to enforce a minimum number of SNPs per window (e.g., at least 10) to ensure stable EHH estimation. Normalization and allele-frequency binning further improve robustness; for sparse datasets, increasing window size (e.g., `--winsize` 200000) and reducing the number of quantile bins (e.g., `--qbins 10`) increases the number of loci per bin and stabilizes normalized estimates. In general, parameter tuning should aim to retain as many usable SNPs as possible while avoiding windows or bins dominated by rare alleles or missing data.

#### 2.6.2 When to use which methods

To choose which method is appropriate for the data we need to consider many important factors and limitation of the dataset available on hand, namely (i) sweep completeness, sample size, and the availability of a suitable reference population, (ii) whether selection is hard or soft, (iii) whether haplotypes are reliably phased, and (iv) whether recombination rate is variable and in such case if recombination distances can be measured accurately using a genetic map. In Figure 1 we show both hard and soft sweep cases for ongoing and completed sweeps, to illustrate why different cases require different methods.

**Figure 1:**
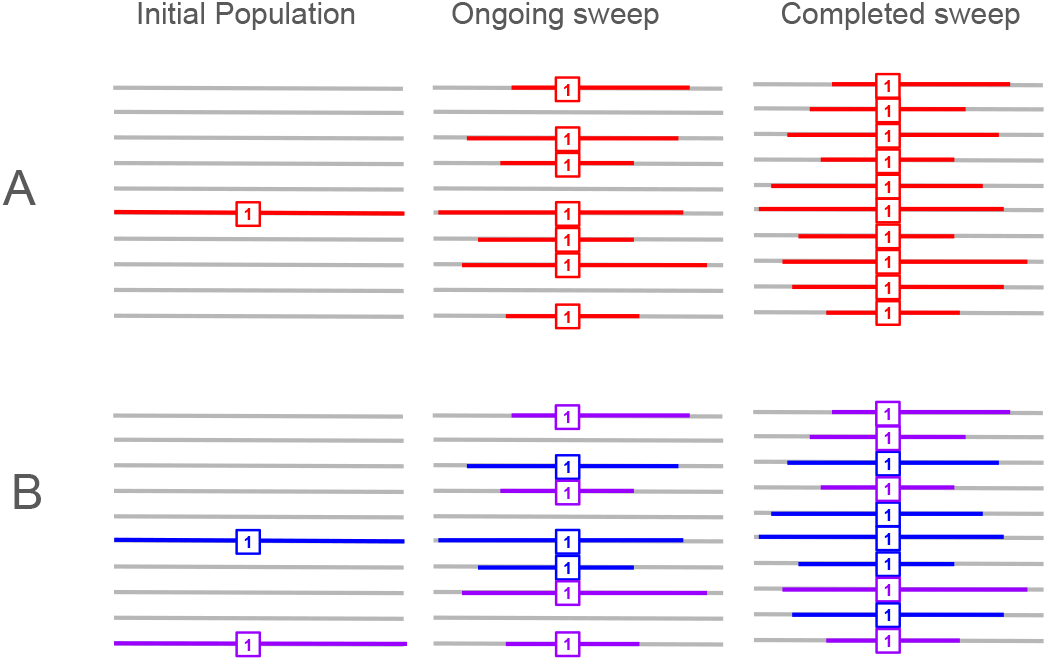
Schematic illustration of haplotype patterns under hard and soft selective sweeps. Panel (A) shows a hard sweep, where a single sweeping haplotype (red) rises from low frequency in the initial population (left column), to an ongoing sweep at ∼70% frequency (middle column), and ultimately to fixation at ∼100% frequency (right column), against a linked haplotype background (gray). Panel (B) shows a soft sweep, where selection acts on multiple haplotype backgrounds (blue and purple), which increase in frequency from the initial population (left column), to ∼70% during the ongoing sweep (middle column), and to near fixation/fixation in the final population (right column), while background haplotypes (gray) are reduced.

For *incomplete sweeps* where the selected allele is segregating at intermediate-to-high frequency, single-population statistics such as iHS or nSL can be used and do not require a reference population when sample size is large enough. An incomplete sweep with the selected allele at ≈70% frequency can be detected using iHS/nSL in a sample of 10 diploid individuals. However, if sampling is reduced, within-population haplotype homozygosity estimates become noisier and iHS/nSL signals may weaken; in such cases, if a suitable reference population is available, XP-based comparisons (XP-EHH/XP-nSL) may retain greater power. In addition, for *completed or near-completed sweeps* (selected allele fixed or nearly fixed), iHS and nSL lose power and may become undefined because the locus may no longer be polymorphic in the sample. In this case, there will be few if any haplotypes lacking the adaptive allele to compare against. Therefore, cross-population methods (XP-EHH or XP-nSL) should improve power whenever a reference population exists.

Sweep type also affects method choice: *hard sweeps* typically generate a single dominant long haplotype and are relatively straightforward to detect with iHS/nSL, whereas *soft sweeps* involve multiple sweeping haplotype backgrounds and may not produce a single extreme haplotype, reducing sensitivity of iHS/nSL. In such cases, iHH12 provides a useful complement because it integrates EHH after combining the two most frequent haplotypes, improving sensitivity to sweeps occurring on multiple backgrounds; as illustrated in Figure 1, when haplotypes from both backgrounds (e.g., the blue and purple haplotypes) rise to high frequency, neither produces a uniquely dominant long-haplotype signal, and iHS/nSL for the derived allele will therefore be weaker. This occurs because extended haplotype homozygosity is effectively split across distinct sweeping backgrounds: in iHS each background is evaluated separately, and because the blue and purple haplotypes are not treated as the same haplotype, the EHH signal is reduced relative to a hard sweep where one dominant haplotype drives the signal.

When phasing is unavailable or uncertain, *unphased* implementations can be used, which is particularly relevant for non-model organisms; however, unphased methods generally require larger sample sizes to maintain power because heterozygous genotypes are effectively not used in the construction of unphased representations, reducing the amount of informative data contributing to the statistic.

*Genetic map quality* affects the reliability of statistics that integrate EHH over genetic distance, i.e., iHS and XP-EHH. When a dense and accurate recombination map is available, these methods can leverage genetic distance to account for local recombination-rate variation. However, in non-model organisms genetic maps might not be available and, when available, are often sparse or heavily interpolated, while recombination-rate variation can be extreme. These issues can distort integration distances (e.g., producing artificially short or collapsed genetic intervals) and increase missing values. In such cases, nSL and XP-nSL can be used as they are less sensitive to map artifacts and recombination-map uncertainty.

For readers interested in exploring the detailed effects of sample size, SNP density, recombination-rate variation, demography and other factors on the power and reliability of these haplotype-based statistics, we suggest consulting the original papers introducing iHS (Voight et al., 2006), nSL (Ferrer-Admetlla et al., 2014), iHH12 (Torres et al., 2018), XP-EHH (Sabeti et al., 2007), XP-nSL (Szpiech et al., 2021), and their unphased versions (Szpiech, 2024). For example, (Ferrer-Admetlla et al., 2014) specifically addresses the effect of recombination rate variation, showing that iHS is more sensitive to local differences in recombination rate, whereas nSL remains comparatively robust under such conditions. It also demonstrates effect of demographic effects such as bottlenecks or expansion in population size.

Finally, several studies have shown that the correlation between iHS and nSL is often modest (Ma et al., 2015), suggesting that these statistics are powered in different regions of parameter space and capture complementary aspects of haplotype structure. This observation has motivated composite approaches, which integrate multiple selection statistics to improve robustness and localization power (Grossman et al., 2013; Randhawa et al., 2014). More generally, because summary statistics vary in informativeness across selective and demographic scenarios, combining them improves reliability; this has been pursued using probabilistic frameworks that model dependencies among statistics (e.g., averaged one-dependence estimators (Sugden et al., 2018)) as well as convolutional neural networks that learn patterns directly from summary statistic representations (Kern and Schrider, 2018)

Overall, low correlation among statistics implies that they often highlight different candidate regions; therefore, relying on a single extreme outlier can be misleading, and candidate regions are best prioritized when supported by multiple complementary signals. `selscan` facilitates this by enabling multiple statistics to be computed efficiently in a single workflow.

#### 2.6.3 Additional options in selscan

`selscan` has many addition options for user convenience. For example, if a genetic map is unavailable but required for iHS and XPEHH, the flag `--pmap` can be set so `selscan` uses physical distances instead. Legacy formats for genetic data are supported, such as TPED (Purcell et al., 2007), HAP (Baumdicker et al., 2022), and transposed HAP (Howie et al., 2009). Additionally, multiple runs of `selscan` can be executed with different parameter configurations and multiple statistics using the --multi-param option. For a full listing and explanation of additional user options, see the `selscan` manual at https://github.com/szpiech/selscan.

### 2.7 Simulated and Empirical Datasets

#### 2.7.1 Simulated Data

Two simple demographic history models, a single population model and a two population divergence model, were simulated to illustrate how the EHH-based statistics implemented in `selscan v3.0` function. For each model, a single hard sweep was simulated, and this was performed both in the presence of background selection and on a purely neutral background. This created four scenarios in total: a hard sweep in a single population, a hard sweep in a single population with background selection, two populations with a hard sweep in one, and two populations with background selection with a hard sweep in one. All simulations were generated using `SLiMv4.2.2` (Haller et al., 2025) with a chromosome length of 10 Mb, a recombination rate of 1 *×* 10^−8^, and a mutation rate 1.29 *×* 10^−8^. In the single population model, the effective population size was set to *N*_*e*_ = 10,000, and, after a 100, 000 generation burn-in, a beneficial mutation (*s* = 0.1) was introduced at position 5 Mb in the 100, 001st generation. The simulation ran for an additional 2, 000 generations until the beneficial mutation reached ∼ 75% frequency. If the beneficial mutation was lost or did not reach ∼ 75% frequency, the simulation was restarted at the 100, 001st generation. At the end of a successful simulation, a VCF file was output. For the scenario with background selection deleterious mutations was introduced at a rate of 9 : 1, neutral:deleterious, with selection coefficients drawn from a gamma distribution with mean 0.1 and shape 0.2. Data for normalization was generated in the same way but without introducing a beneficial allele. 100 neutral replicates were generated for each scenario (with and without background selection). The neutral replicates were used to normalize the replicate containing the selected allele. Fifty individuals were sampled for each simulation. The resulting variants were filtered for biallelic SNPs and removed if the minor allele frequency below 0.05.

In the two population model, an ancestral population with effective population size *N*_*e*_ = 10,000 was simulated for a 100, 000 generation burn-in. At generation 100, 500, the ancestral population split into two populations each with effective population size *N*_*e*_ = 10,000. At generation 102, 000, a beneficial mutation (*s* = 0.1) was introduced into one population at position 5 Mb. The simulation ran for an additional 2, 000 generations until the beneficial mutation reached ∼ 75% frequency. If the beneficial mutation was lost or did not reach ∼ 75% frequency, the simulation was restarted at the 100, 001st generation. At the end of a successful simulation, a VCF file was output. For the scenario with background selection deleterious mutations was introduced at a rate of 9 : 1, neutral:deleterious, with selection coefficients drawn from a gamma distribution with mean 0.1 and shape 0.2. Data for normalization was generated in the same way but without introducing a beneficial allele. For each scenario, 100 neutral replicates were generated (with and without background selection). The neutral replicates were used to normalize the replicate containing the selected allele. Fifty individuals were sampled for each simulation. The resulting variants were filtered for biallelic SNPs.

For the two population model, with and without background selection, we investigated the effects of uneven sample sizes between the two populations. We tested sampling ten individuals from the first population and fifty individuals from the second population, and we tested sampling fifty individuals from the first population and ten individuals from the second population. This was done for the replicates containing the sweep as well as all neutral replicates. For each scenario, the resulting neutral replicates were used to normalize the replicate containing the sweep.

#### 2.7.2 Empirical Data

The phased and imputed genomes of two populations from the 1000 Genomes Project were used: 99 Utah individuals with Northern and Western European ancestry (CEU) and 108 Yoruba individuals from Ibadan, Nigeria (YRI) (Consortium et al., 2015; Byrska-Bishop et al., 2022). All autosomes were extracted and were restricted to biallelic SNPs. For iHS, nSL, and iHH12 calculations, selscan filters sites with a minor allele frequency less than 0.05.

Gene annotations were obtained from the GENCODE database (GRCh38) using the GTF file from (Frankish et al., 2021).

## 3 Results

In this section, we summarize the outcomes of applying `selscan`, and subsequently, `selscan norm` to both simulated and empirical datasets.

### 3.1 Simulated Data

#### 3.1.1 Importance of Normalization

Figures 2 and 3 illustrate the importance of normalization for iHS and nSL. In Figures 2a and 3a, the raw statistic from the replicate containing the hard sweep is plotted against derived allele frequency, showing a clear correlation. This is because iHS and nSL are log-ratios of *EHH*_*c*_ (Equation 3), which is summarizing haplotype lengths surrounding either the ancestral or derived alleles. As low-frequency alleles are likely to be younger and therefore segregate on longer haplotypes, even under neutrality, this implies larger log-ratio scores for low frequency alleles. Figures 2b and 3b show how normalization in frequency bins (Equation 13) corrects for this frequency bias. To underscore the importance of normalization, Figures 2c and 3c contrast raw |*scores*| and normalized |*scores*| as a function of genome location for the sweep simulation, clearly demonstrating that without normalization the sweep signal would be lost.

**Figure 2:**
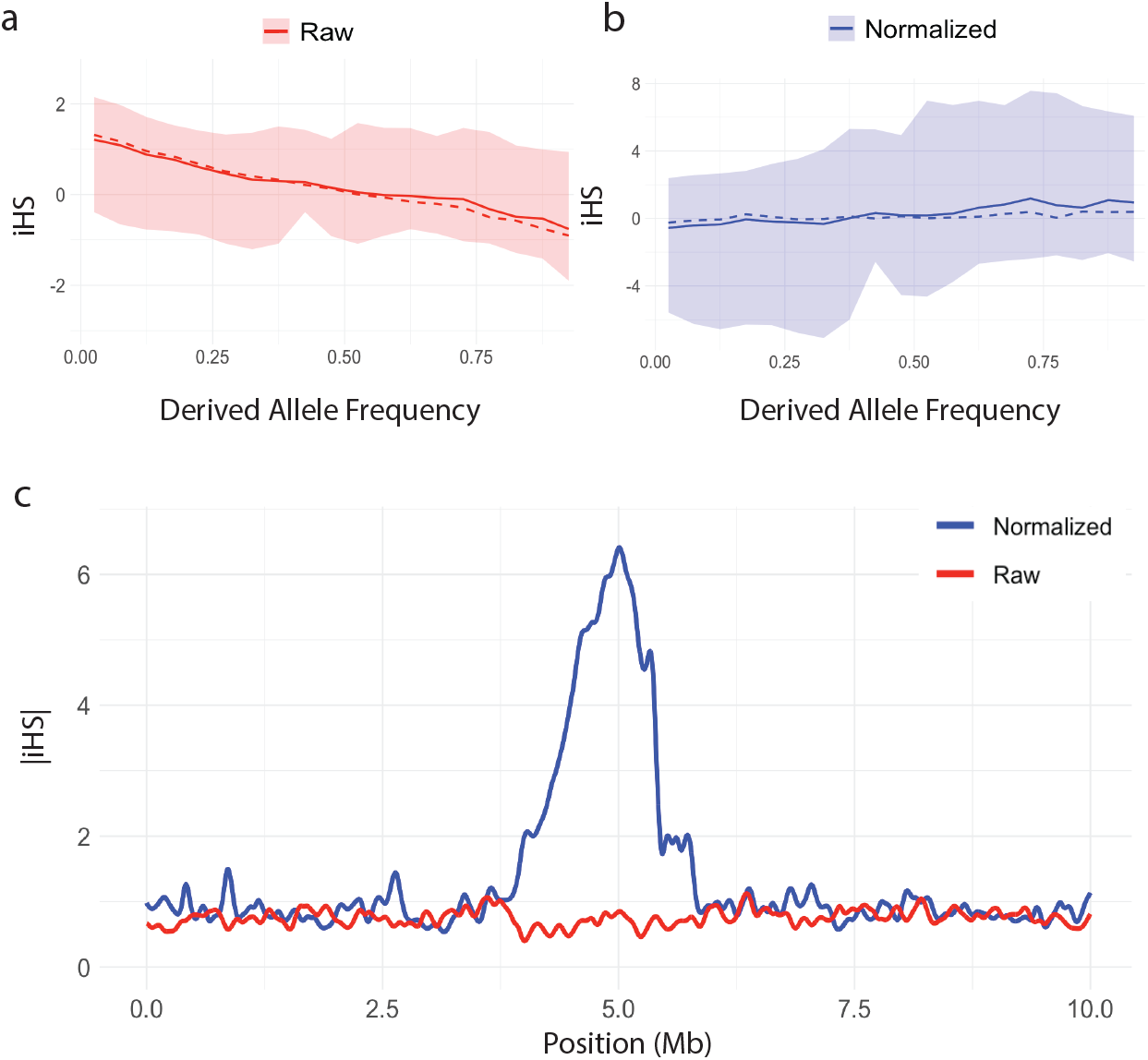
The mean (solid line), median (dashed line), and 1st–99th percentile range (shaded region) for **(a)** raw iHS and **(b)** normalized iHS statistic plotted against derived allele frequency. **(c)** Spline smoothed raw and normalized curves of |*iHS*| in the vicinity of a simulated hard sweep at 5Mb.

**Figure 3:**
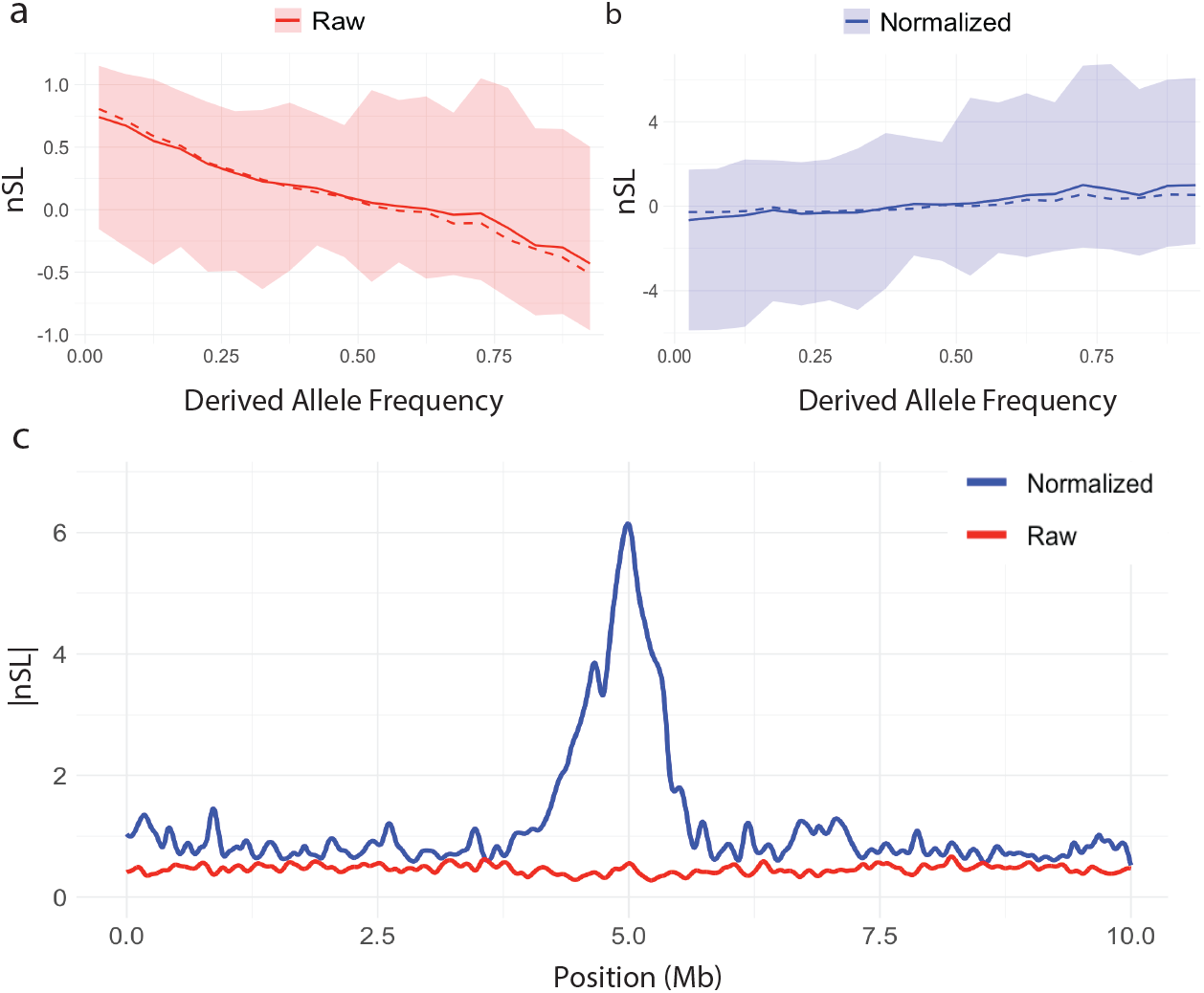
The mean (solid line), median (dashed line), and 1st–99th percentile range (shaded region) for **(a)** raw nSL and **(b)** normalized nSL statistic plotted against derived allele frequency. **(c)** Spline smoothed raw and normalized curves of |*nSL*| in the vicinity of a simulated hard sweep at 5Mb.

XPEHH and XPnSL, however, do not suffer from this dependence on derived allele frequency, as they are comprised of the *EHH* (Equation 1) of all haplotypes within each population, instead of restricting to haplotypes carrying a particular allele. This is illustrated in Figure 4. We show similar behavior for iHH12 in Figure 5. However, normalization is still recommended, as it puts different experiments on the same scale and allows for easy downstream statistical analyses which may assume variables are distributed Normally. We emphasize the that two-population statistic normalization does not depend on frequency (Equation 14) unlike single-population statistics.

**Figure 4:**
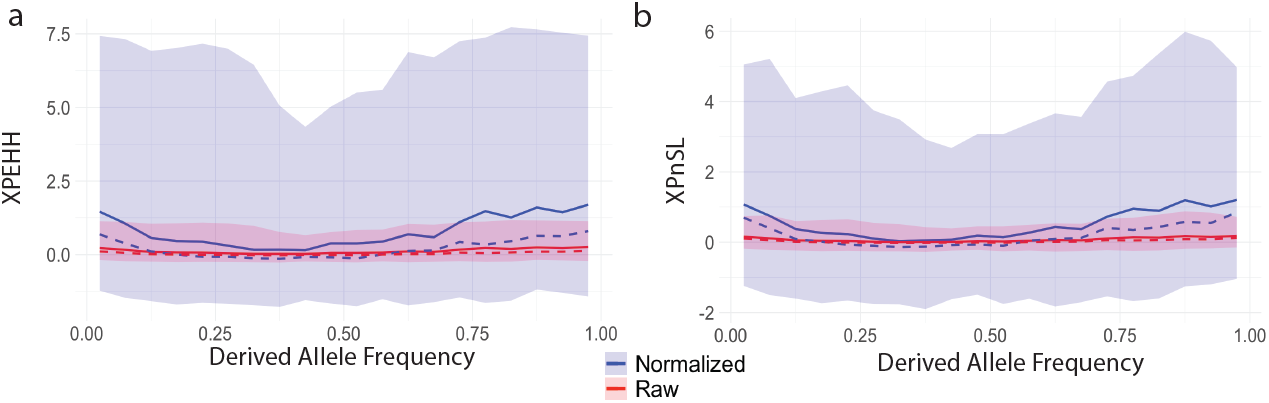
The mean (solid line), median (dashed line), and 1st–99th percentile range (shaded region) for **(a)** raw (red) and normalized (blue) XPEHH and **(b)** raw (red) and normalized (blue) XPnSL statistics plotted against derived allele frequency.

**Figure 5:**
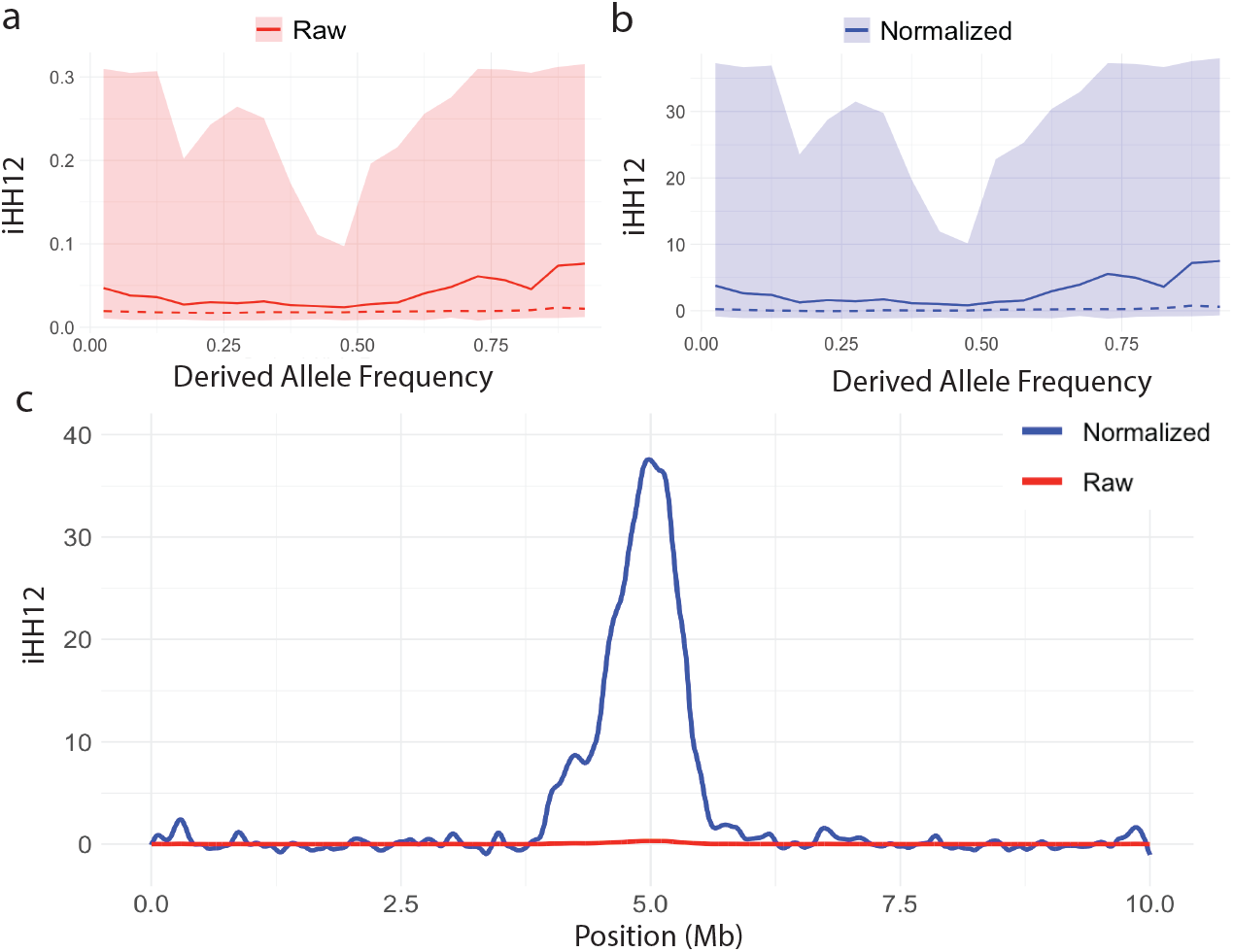
The mean (solid line), median (dashed line), and 1st–99th percentile range (shaded region) for **(a)** raw iHH12 and **(b)** normalized iHH12 statistic plotted against derived allele frequency. **(c)** Spline smoothed raw and normalized curves of |*iHH*12| in the vicinity of a simulated hard sweep at 5Mb.

#### 3.1.2 Single-population Statistics

Figure 6 illustrates how EHH can distinguish between genome regions with an adaptive allele sweeping and genome regions without an adaptive allele. Figure 6a plots the *EHH* of the entire sample in the vicinity of a sweep (dotted line), which decays slowly over distance, in contrast to the *EHH* in a neutral region (solid line) which decays quickly. Figure 6b illustrates how computing *EHH*_*c*_ can distinguish between the sweeping haplotype and the non sweeping haplotype. In the sweep simulation, *EHH*_1_ is computed among the haplotypes carrying the adaptive derived allele, and haplotype homozygosity decays slowly. However, in the same simulation at the same locus, *EHH*_0_ is computed among the haplotypes not carrying the adaptive allele, and haplotype homozygosity decays quickly. Whereas in the neutral simulation both *EHH*_1_ and *EHH*_0_ decay quickly as neither set of hapotypes contain an adaptive allele. Figures 6c and Figure 6d show similar patterns in the context of background selection. Based on these patterns, iHH12, iHS, and nSL are designed to efficiently summarize the contrasts in these curves for large-scale querying of millions of loci across a genome.

**Figure 6:**
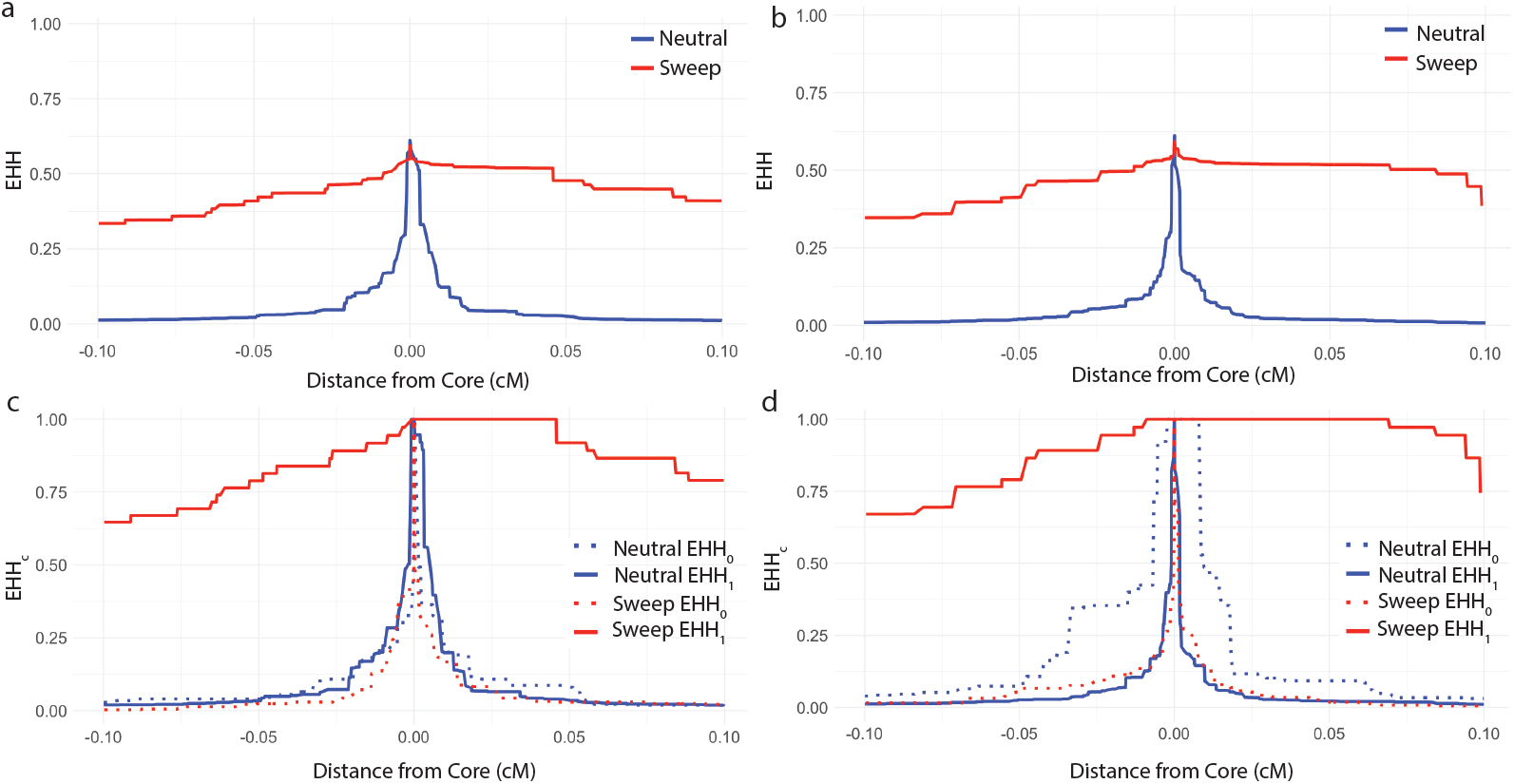
**(a)** *EHH* curves for a simulated hard sweep and a simulated neutral region. **(b)** *EHH*_*c*_ curves for each of the core haplotypes (0 and 1) for a simulated hard sweep and a simulated neutral region. **(c)** *EHH* curves for a simulated hard sweep and a simulated region without a sweep in the presence of background selection. **(d)** *EHH*_*c*_ curves for each of the core haplotypes (0 and 1) for a simulated hard sweep and a simulated region without a sweep in the presence of background selection.

Figure 7 illustrates how iHH12 (Figure 7a), iHS (Figure 7b), and nSL (Figure 7c) can identify a sweep in the center of a simulated 10Mb region. For all three statistics, regardless of the presence of background selection and regardless of whether allele phase is utilized (iHS and nSL only), a clear peak forms in the vicinity of where the adaptive allele was placed. This suggests that the EHH-based single-population statistics are robust to background selection.

**Figure 7:**
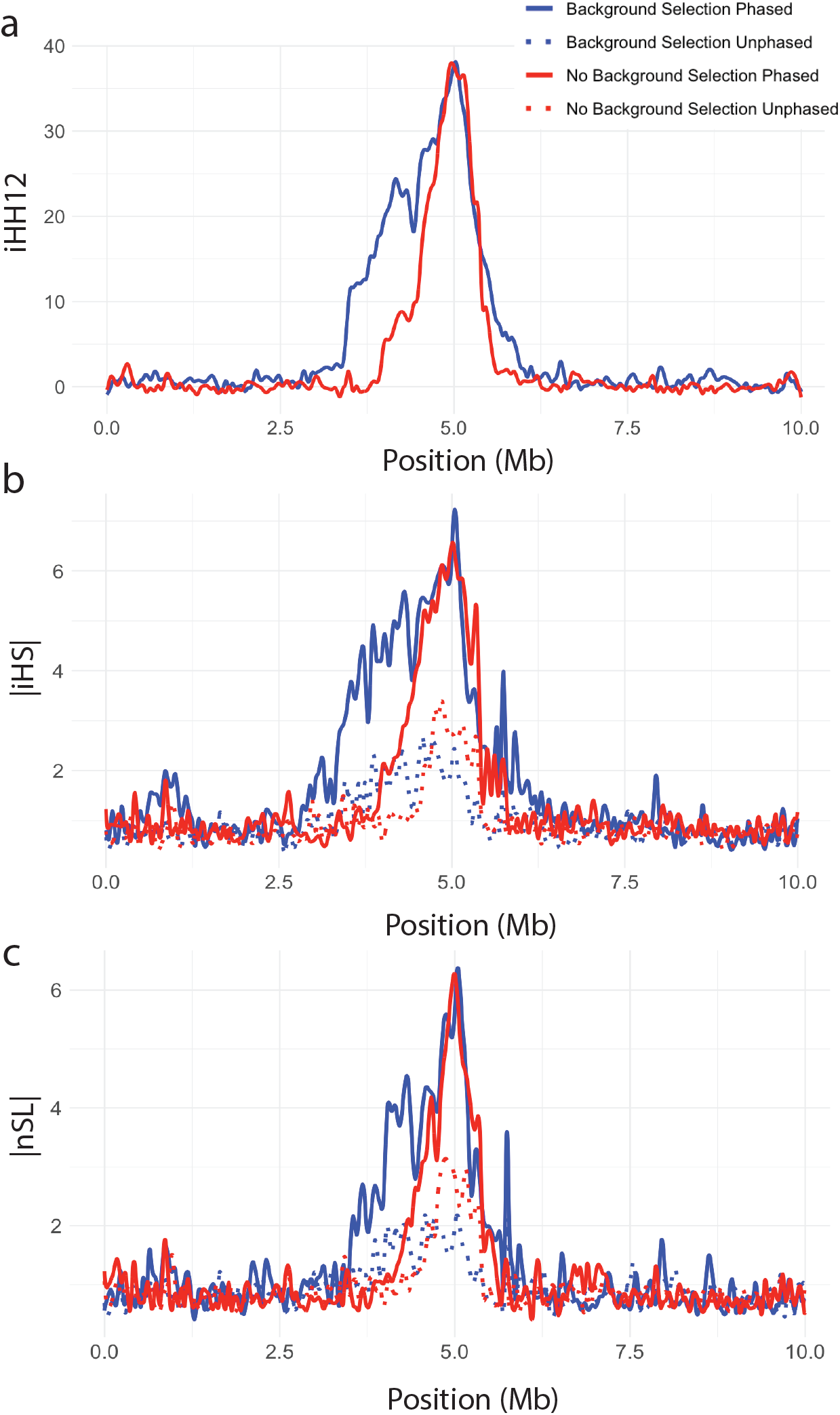
Normalized **(a)** *iHH*12, **(b)** |*iHS*|, and **(c)** |*nSL*| plotted against genomic position for a simulated hard sweep with and without background selection. All curves were splined smoothed.

#### 3.1.3 Two-population Statistics

The two-population statistics XPEHH and XPnSL are designed to summarize and contrast EHH patterns between two closely related populations to identify local adaptation. Figure 8 illustrates how these two statistics peak in the center of the simulated region where the adaptive allele was place. Figures 8a plot XPnSL and Figures 8b plot XPEHH. In addition to testing the influence of phased genotypes and background selection, the influence of unequal sample sizes among the two populations was considered, showing minimal influence 9. This suggests that the EHH-based two-population statistics are robust to background selection and uneven samples size.

**Figure 8:**
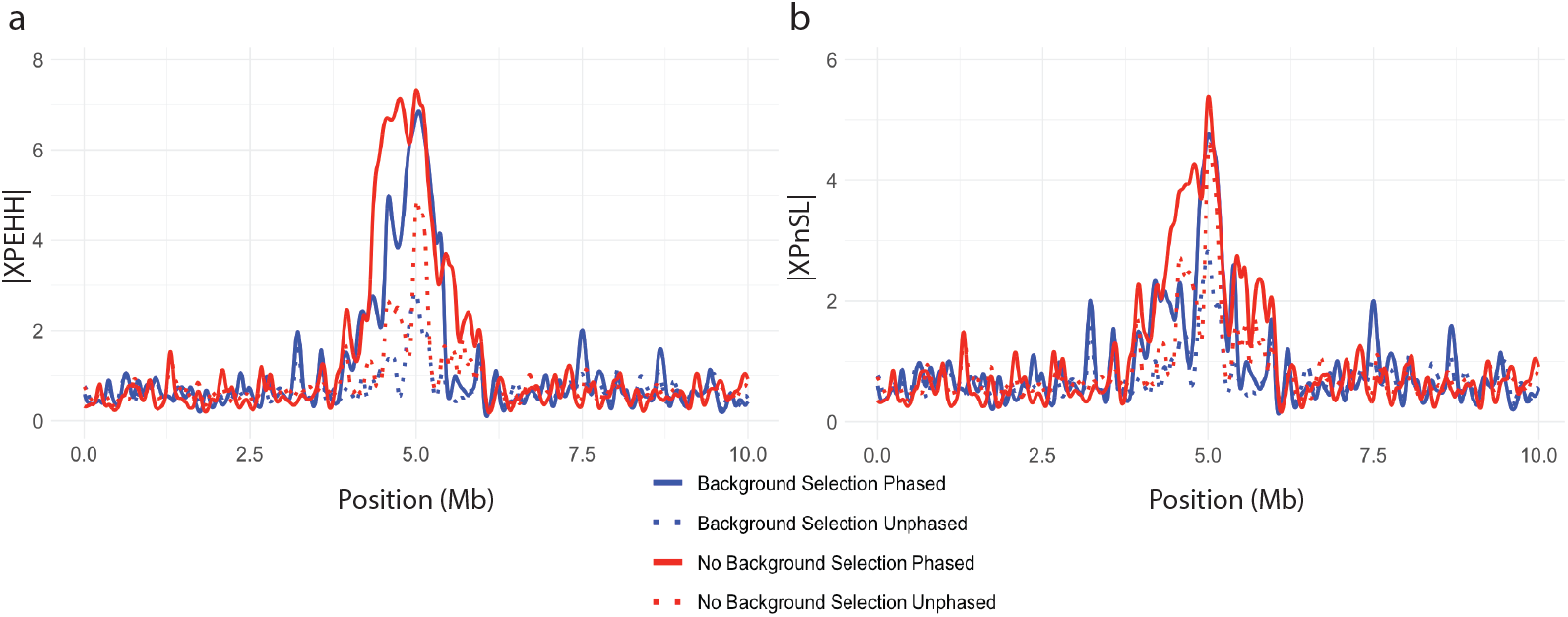
Normalized **(a)** XPEHH and **(b)** XPnSL plotted against genomic position for a simulated hard sweep with and without background selection. All curves were splined smoothed.

### 3.2 Basic downstream analysis with selscan

Section 3.1 demonstrates how `selscan`‘s EHH-based statistics behave under simulated scenarios. However, `selscan`‘s `norm` subcommand contains basic downstream analysis tools that are best illustrated using an empirical dataset.

This section illustrates an example of `selscan norm`‘s window-based outlier detection and gene annotation functions can be used to find genes putatively under positive selection. This example uses the European CEU and Yoruba YRI populations from the 1000 Genomes Project (Consortium et al., 2015; Byrska-Bishop et al., 2022) and focuses on the well-known selection signal in the LCT/MCM6 region of the human genome. It involves lactase persistence, a phenotype which has evolved multiple times in humans (Tishkoff et al., 2007). In the CEU population, this phenotype is caused by a variant within an intron in the MCM6 gene upstream of LCT, allowing lactase expression into adulthood (Ingram et al., 2009).

`selscan` was used to compute iHS, nSL, iHH12 across all autosomes in the CEU population, and XPEHH and XPnSL were computed on all autosomes comparing CEU and YRI populations. Genome-wide normalization of all statistics was performed using `selscan norm`, with iHS and nSL being normalized with 100 frequency bins. For iHS, nSL and iHH12 normalization, we used respective scores from all CEU autosomes, and for XPEHH and XPnSL, we used respective scores from all autosomes with CEU as the target population and YRI as the reference population. Figure 10 shows the raw-score distributions (Figure 10a and c) and normalized-score distributions (Figure 10b and d) for scores on chromosome 2.

**Figure 9:**
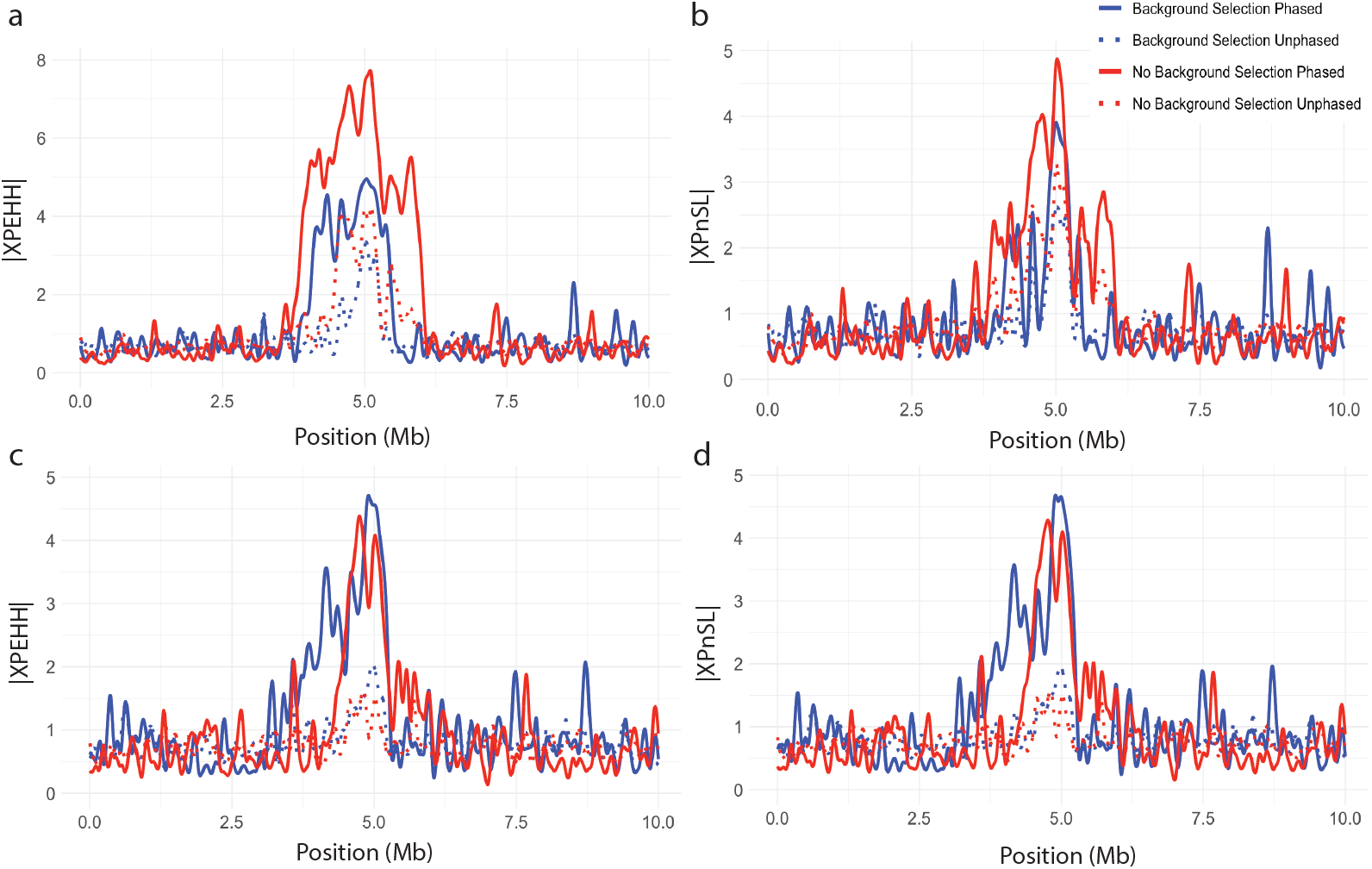
Normalized **(a)** XPEHH with 50 sample from P1 and 10 samples from P2 **(b)** XPnSL with 50 sample from P1 and 10 samples from P2 **(c)** XPEHH with 10 sample from P1 and 50 samples from P2 **(d)** XPnSL with 10 sample from P1 and 50 samples from P2. All are plotted against genomic position for a simulated hard sweep with and without background selection. All curves were splined smoothed.

**Figure 10:**
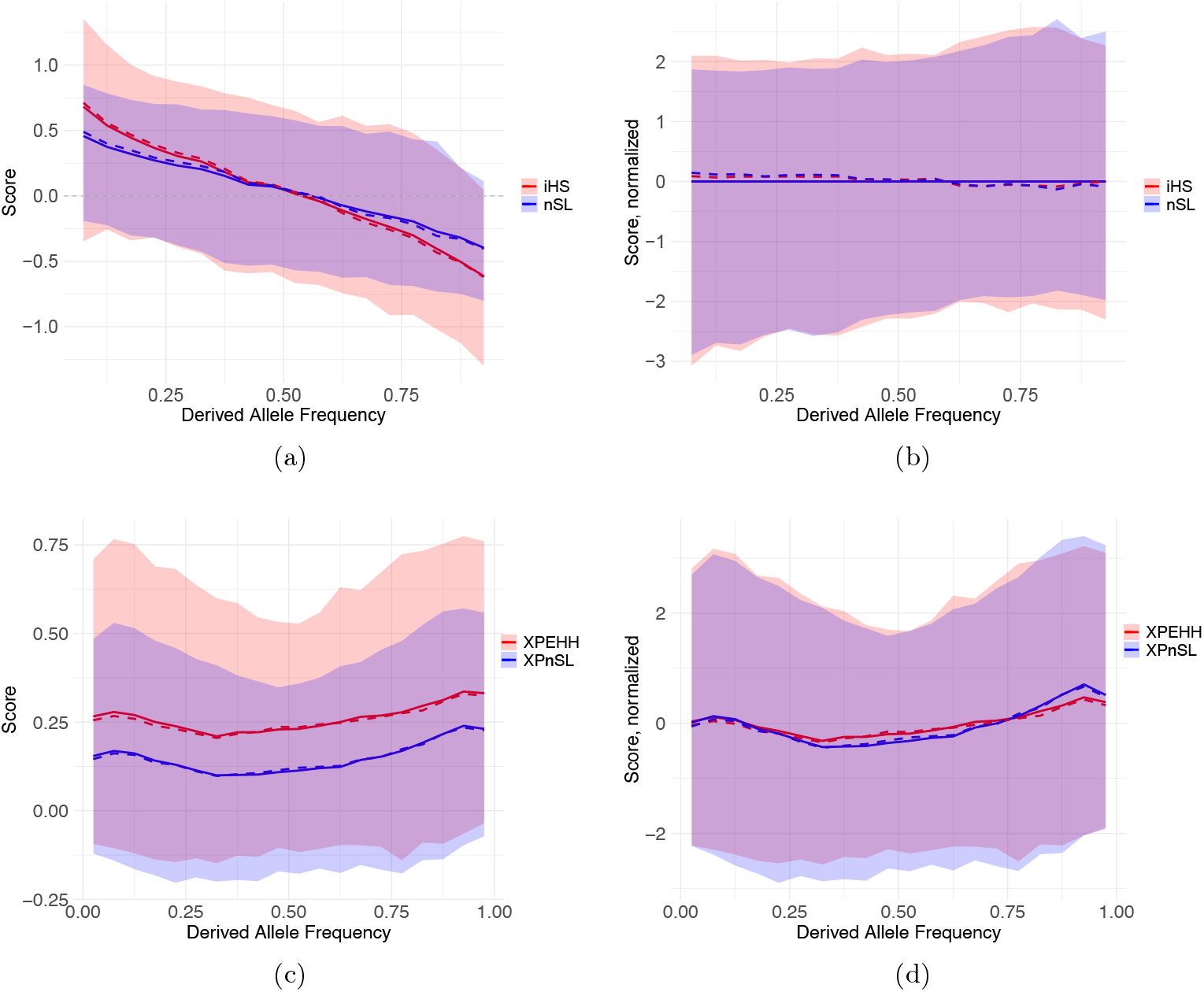
Distribution of EHH-based statistics versus derived allele frequency (DAF). **(a)** raw his (red) and nSL (blue) scores. **(b)** normalized iHS (red) and nSL (blue) scores. **(c)** raw XPEHH (red) and XPnSL (blue) scores. **(d)** normalized XPEHH (red) and XPnSL (blue) scores. All panels show the mean (solid line), median (dashed line), and 1st–99th percentile range (shaded area).

#### 3.2.1 Analysis of chromosome 2 identifies the LCT/MCM6 locus

Focusing on chromosome 2 XPEHH scores, `selscan norm` was used to identify window-based outliers (default parameters) and annotated with overlapping gene regions. Table 1 shows `selscan norm` output restricted to windows falling in the top 1% and sorted by maximum observed score per window. As anticipated, the LCT/MCM6 locus is found at the very top of this list, confirming that this locus is among the strongest selected on chromosome 2.

**Table 1:** Top 20 windows on chromosome 2 from the top 1% of XP-EHH scores (sorted by maximum score column) showing the maximum score per window and the genes overlapping each window.

| Start (bp) | End (bp) | nSnps | Frac of SNPs<br>with score<br>above threshold | Maximum<br>Score | Overlapped<br>Genes |
| --- | --- | --- | --- | --- | --- |
| 135,800,001 | 135,900,000 | 1,946 | 1.00 | 6.19 | **LCT, MCM6** |
| 135,700,001 | 135,800,000 | 1,838 | 1.00 | 5.86 | R3HDM1,<br>UBXN4, **LCT** |
| 135,900,001 | 136,000,000 | 1,800 | 1.00 | 5.70 | DARS1 |
| 135,400,001 | 135,500,000 | 2,288 | 1.00 | 5.51 | ZRANB3 |
| 135,200,001 | 135,300,000 | 1,958 | 1.00 | 5.16 | ZRANB3 |
| 135,300,001 | 135,400,000 | 2,016 | 1.00 | 5.10 | ZRANB3 |
| 135,000,001 | 135,100,000 | 1,900 | 1.00 | 4.99 | MAP3K19,<br>RAB3GAP1 |
| 135,100,001 | 135,200,000 | 1,752 | 1.00 | 4.88 | RAB3GAP1,<br>ZRANB3 |
| 135,500,001 | 135,600,000 | 1,800 | 1.00 | 4.81 | ZRANB3,<br>R3HDM1 |
| 236,200,001 | 236,300,000 | 2,095 | 0.51 | 4.59 | ASB18 |
| 237,500,001 | 237,600,000 | 2,822 | 0.56 | 4.29 | MLPH, PRLH,<br>RAB17 |
| 136,500,001 | 136,600,000 | 2,245 | 0.93 | 4.26 | - |
| 134,900,001 | 135,000,000 | 2,015 | 0.83 | 4.15 | ACMSD,<br>CCNT2,<br>MAP3K19 |
| 237,400,001 | 237,500,000 | 2,309 | 0.72 | 4.07 | COL6A3, MLPH |
| 134,800,001 | 134,900,000 | 1,991 | 1.00 | 4.04 | ACMSD |
| 6,100,001 | 6,200,000 | 2,684 | 0.59 | 4.00 | - |
| 177,600,001 | 177,700,000 | 2,215 | 0.52 | 3.93 | IFT70A,<br>PDE11A |
| 136,400,001 | 136,500,000 | 2,186 | 0.89 | 3.78 | - |
| 126,800,001 | 126,900,000 | 2,447 | 0.40 | 3.63 | TEX51 |
| 236,500,001 | 236,600,000 | 2,361 | 0.90 | 3.61 | IQCA1, ACKR3 |

Zooming into the LCT/MCM6 locus, Figure 11a plots normalized |*iHS*| as a function of genome position on chromosome 2, and Figure 11b plots normalized *XPEHH* scores, as a function of genome position on chromosome 2. Both plots show broad peaks overlapping the LCT/MCM6, suggesting a clear signal of positive selection at this locus.

**Figure 11:**
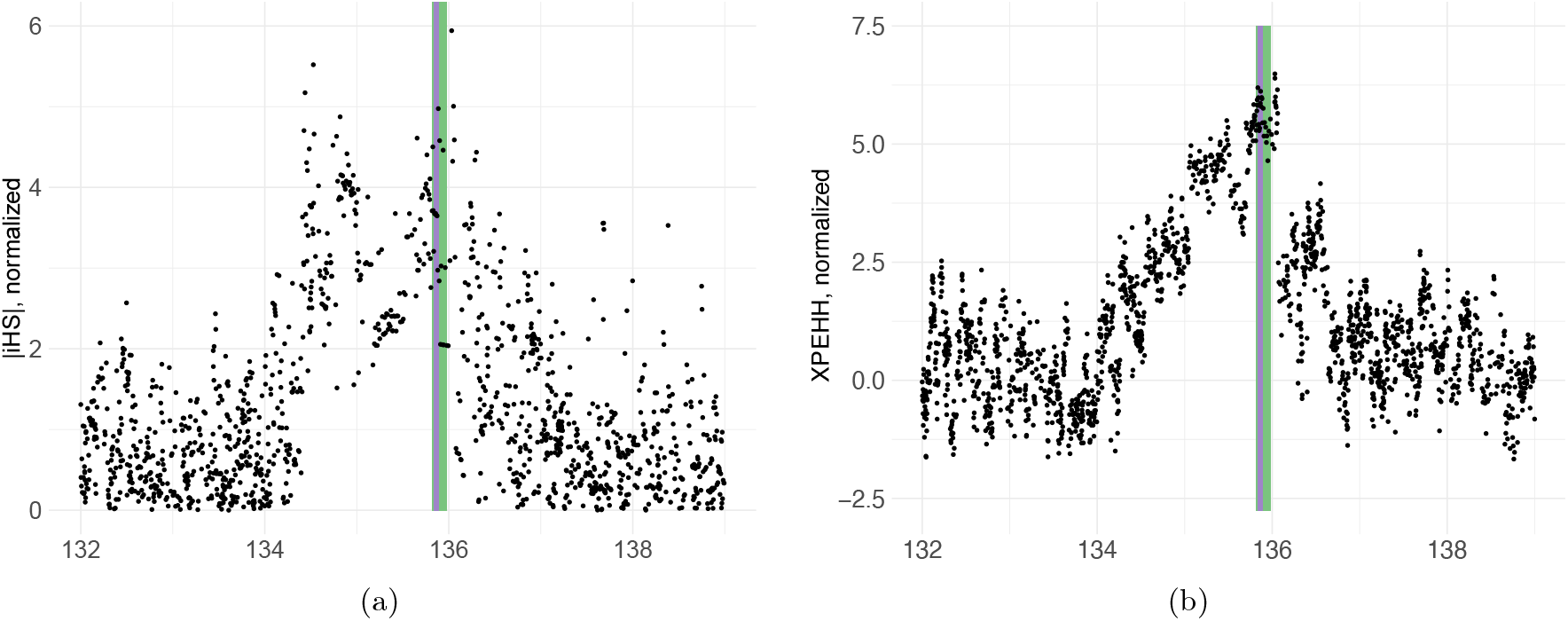
Normalized **(a)** |*iHS*| plotted against chromosome 2 position in CEU, and normalized XPEHH between CEU and YRI plotted against chromosome 2 position. The green shaded area highlights LCT, and the purple shaded area highlights MCM6.

`selscan norm` also creates a gene table (Table 2) for each gene. This table provides information on a per-gene basis including gene length (as given in the BED file), number of overlapping windows, max observed score in the gene, and an adjusted max observed score. The latter is provided to account for varying gene length. Figure 12a shows the correlation of XPEHH score with gene length, and Figure 12b shows the relationship after regressing out its influence. Table 2 is sorted by length-corrected maximum observed score, and LCT and MCM6 fall at the very top.

**Table 2:** The table shows gene-level output from chromosome 2 for the CEU vs YRI comparison. Normalized XP-EHH scores were computed at the SNP level and standardized using the distribution across all autosomes. For each gene, all SNPs falling within the gene boundaries were collected. SNPs with normalized XP-EHH values greater than 2 were considered critical, and the “Fraction of Critical SNPs” column reports the proportion of such SNPs within each gene. Gene-level statistics were computed, including the maximum score, defined as the highest normalized XP-EHH value among all SNPs within the gene boundaries, and the length-corrected maximum score, which adjusts this maximum based on the distribution of scores across all genes on all autosomes to account for gene length. LCT and the gene MCM6, which contains an enhancer region approximately 14 kb upstream of LCT that regulates lactase expression, emerged as the top candidates on chromosome 2. We report the top 20 highest-scoring genes ranked by their length-corrected maximum score.

| Gene Name | Length (bp) | # SNPs | Fraction of Critical SNPs | Max Score | Length-Corrected Max Score |
| --- | --- | --- | --- | --- | --- |
| **MCM6** | 36,854 | 269 | 1.00 | 6.3 | 5.2 |
| **LCT** | 49,335 | 367 | 1.00 | 6.4 | 5.2 |
| UBXN4 | 43,433 | 299 | 1.00 | 5.8 | 4.6 |
| DARS1 | 80,220 | 468 | 1.00 | 5.9 | 4.5 |
| R3HDM1 | 193,867 | 1,145 | 1.00 | 6.0 | 4.3 |
| ZRANB3 | 394,313 | 2,856 | 1.00 | 5.7 | 3.7 |
| RAB3GAP1 | 124,385 | 816 | 1.00 | 5.2 | 3.6 |
| ANKRD36B | 97,215 | 410 | 0.37 | 4.6 | 3.1 |
| MLPH | 69,895 | 952 | 0.92 | 4.4 | 3.1 |
| POTEI | 50,253 | 382 | 0.52 | 4.2 | 2.9 |
| MAP3K19 | 82,984 | 664 | 0.94 | 4.3 | 2.8 |
| ACMSD | 64,595 | 459 | 1.00 | 4.2 | 2.8 |
| ARHGEF4 | 210,350 | 1,982 | 0.15 | 4.5 | 2.8 |
| LRRTM4 | 845,635 | 9,739 | 0.13 | 5.0 | 2.7 |
| GALNT5 | 60,914 | 507 | 0.62 | 4.0 | 2.7 |
| DOK1 | 8,680 | 54 | 1.00 | 3.3 | 2.7 |
| CIB4 | 60,162 | 639 | 0.34 | 3.9 | 2.6 |
| IQCA1 | 183,570 | 1,856 | 0.12 | 4.2 | 2.5 |
| RAB6D | 3,677 | 37 | 0.65 | 2.7 | 2.4 |
| MYO3B | 477,027 | 4,911 | 0.04 | 4.4 | 2.3 |

**Figure 12:**
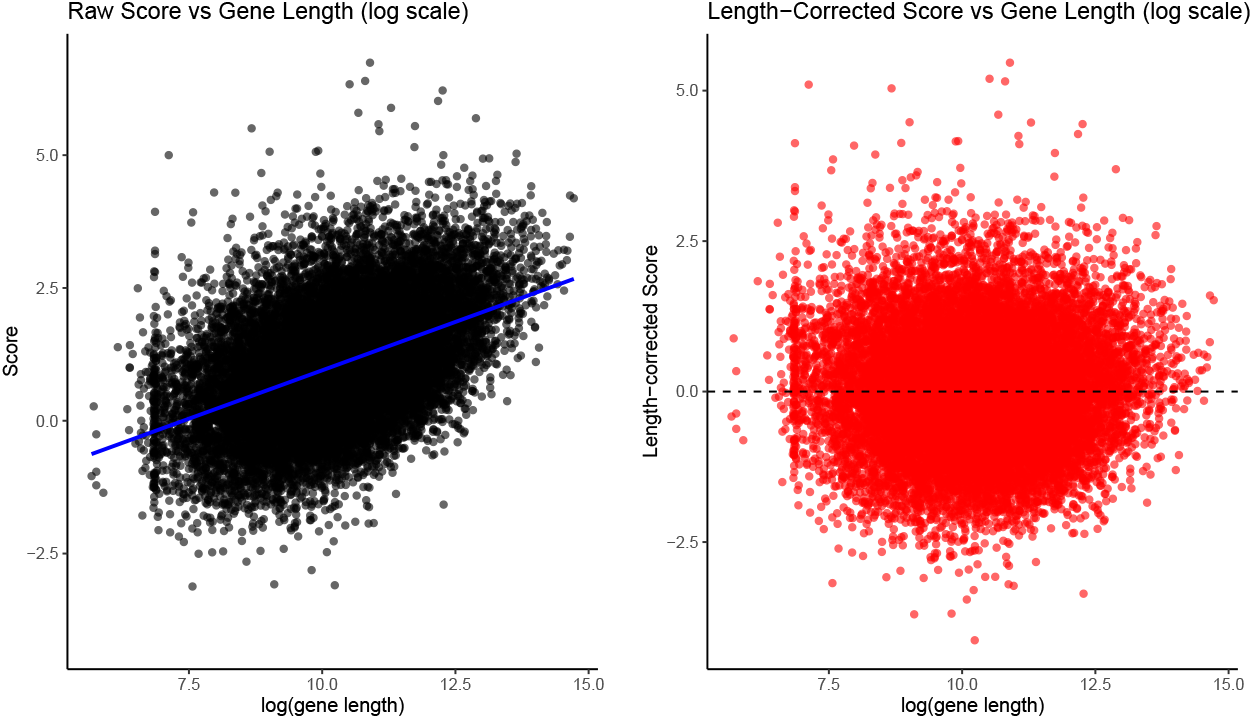
Length correction for XP-EHH scores in chromosome 2. Left panel: raw per-gene scores plotted against log(gene length), with a linear regression trend line (blue) showing the overall relationship. Right panel: length-corrected scores (residuals from the regression) plotted against logarithm of gene length, with a dashed line indicating the mean.

#### 3.2.2 Comparison of different statistics in CEU chr2

Figure 13 and Figure 14 compares selection signals across chromosome 2 for CEU. Each statistic operates in its own parameter space, and all detect the strong LCT–MCM6 signal, even without phase information. In our results, iHS and nSL (phased) show good correlation. At high SNP density (as in our data), these differences are small, but we can expect larger differences that break this correlation in other data where recombination rate varies, as discussed and experimentally shown in the original nSL paper. iHH12 captures soft sweeps and identifies more candidate regions overall, including rare, long haplotypes that iHS and nSL largely miss. The two unphased statistics are more similar to each other, as are the two phased statistics, while comparisons between phased and unphased versions show smaller correlation, reflecting slight information loss due to ignoring heterozygotes.

**Figure 13:**
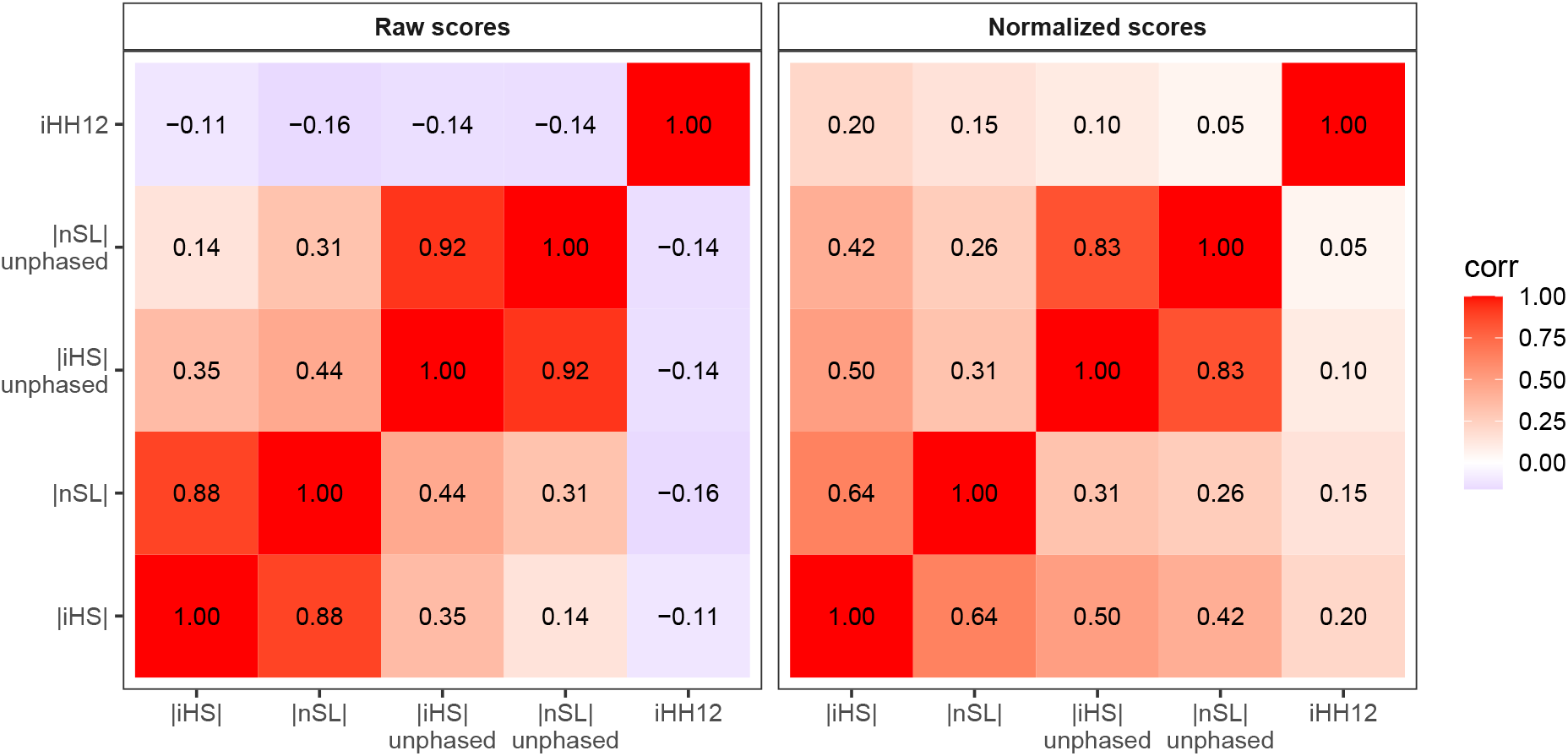
Pearson correlation of five selection statistics across chromosome 2 in the CEU population. Normalized scores (left panel) and raw scores (right panel) are shown for all single population statistics. Only positions where all statistics are defined are included. Values within tiles indicate the pairwise correlation.

**Figure 14:**
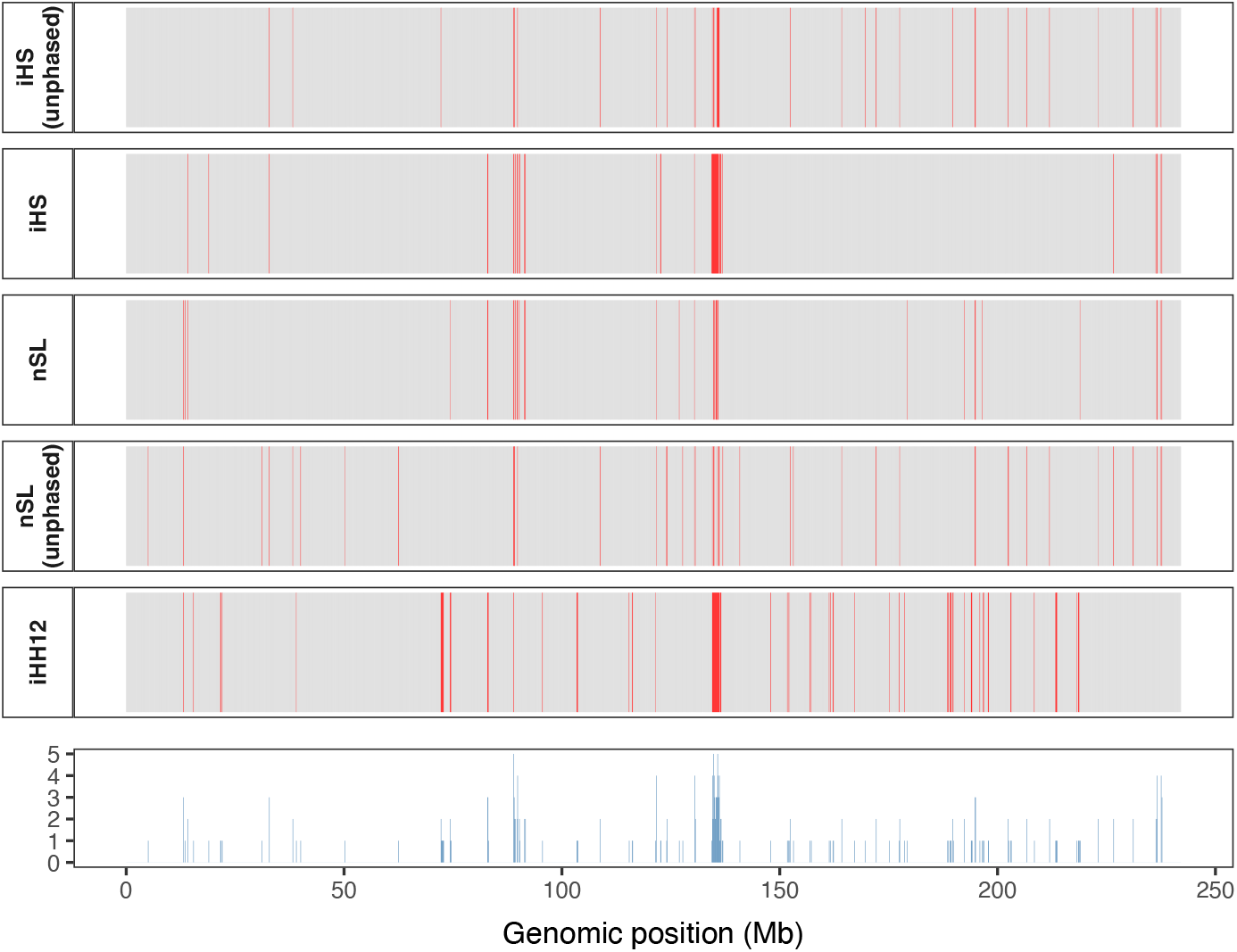
Comparison of aggregated selection scores (maximum per 100 kb window) across chromosome 2 in the CEU population. In the top panel, each tile represents a window for five statistics (iHS, nSL, iHS unphased, nSL unphased, iHH12), with red indicating windows in the top 1% and gray otherwise. The bottom panel shows the consensus score, indicating the number of statistics that are significant in each window (range 0–5). A peak in consensus is observed around the LCT–MCM6 region (≈135–136 Mb), reflecting strong concordance of high-scoring windows in this well-known lactose persistence locus. Only windows where all statistics are defined are included.

## 4 Discussion

The use of selection statistics is ubiquitous in modern evolutionary genomics. The ability to identify putatively selected loci is of great importance in forming our understanding of adaptation of organisms to myriad selection pressures (Fu and Akey, 2013). Therefore, the computation and analysis of selection statistics should be consistent across studies to maintain reproducibility of results and to ensure robust and reliable inference.

Many of the most widely used selection statistics for recombining genomes are based on Expected Haplotype Homozygosity (EHH), which measures the decay of haplotype identity away from a locus of interest (Sabeti et al., 2002, 2007). Here we provide the formal definition of EHH and the formal definitions of the EHH-based statistics iHS, nSL, iHH12, XPEHH, and XPnSL. All statistics and their unphased equivalents are implemented in the software `selscan`. We also introduced the `selscan v3.0` subcommand `norm. selscan norm` is used to perform essential normalization computations, and can also be used for window-based outlier detection and gene-based annotation.

We demonstrated the use of `selscan` on both simulated and empirical genetic data. Under ideal parameters, simulations illustrated the statistics’ ability to identify loci under positive selection in the presence and absence of background selection. This supports the use of statistics based on EHH, since other selection statistics based on the site frequency spectrum can fail to uncover signals of positive selection in this context (but see (Huber et al., 2016; DeGiorgio et al., 2016)). Further, we used `selscan norm` to show how, without normalization, the selection signal is dampened. Finally, we performed a downstream analysis with `selscan norm` on data from the 1000 Genomes Project, focusing on the well-known selection signal at the LCT/MCM6 locus in European individuals (Tishkoff et al., 2007). We showed how the results produced by `selscan` and `selscan norm` quickly produce window-level and gene-level data that can be presented as-is or used in further statistical analyses.

Ultimately, we hope to create a resource for the computation and interpretation of EHH-based selection statistics that is accessible to readers from various backgrounds. While `selscan` is designed to be user friendly, we note that the efficiency of `selscan`‘s core algorithm in conjunction with `selscan norm` offers the rapid computation and analysis of statistics at millions of loci (Rahman et al., 2025). This offers a potential future avenue to scale EHH-based selection statistics to massive empirical and synthetic datasets, and for use in the training of machine learning models (Amin et al., 2024).

## Acknowledgments

Computations for this research were performed using the Pennsylvania State University’s Institute for Computational Data Sciences’ Roar supercomputer. This work was supported by the National Institute of General Medical Sciences of the National Institutes of Health award number R35GM146926, and start-up funds from the Pennsylvania State University’s Department of Biology.

